# Monkeypox Virus Clade IIb Isolate Exhibits Reduced Virulence Relative to Clade IIa Isolates in Multiple Murine Models

**DOI:** 10.64898/2025.12.10.693532

**Authors:** Stephanie V Trefry, Santiago Vidal-Freire, Mayanka Awasthi, Abel D Ordonez, Brett P Eaton, Christy N Raney, Amy L Cregger, Chase A Gonzales, Robert N Enamorado, Nelson A Martinez, Deborah S Gohegan, Matthew G Lackemeyer, Tinoush Moulaei, Natasza E Ziółkowska, Scott J Goebel, Gustavo Palacios, Sina Bavari, Farooq Nasar

## Abstract

Monkeypox virus (MPXV) is the causative agent of mpox disease in humans. The virus is comprised of two clades, Central African clade I and West African clade II with case fatality rates of ∼11 and ∼4%, respectively. Since the discovery of mpox disease in 1970, the virus has been restricted to Africa. However, in 2022, a previously unrecognized subclade IIb caused the largest global outbreak of mpox disease with a case fatality rate of ∼0.2%. The difference in virulence of MPXV subclades in human infection warrants further investigation, however, one critical limitation is the lack of susceptible small animal models. In this study, we investigated the susceptibility of four murine models, including CAST-EiJ and three immunocompromised models (C57BL/6 *Ifnar*^−/−^, C57BL/6 *Ifngr*^−/−^, and C57BL/6 *Ifnar*^−/−^/*Ifngr*^−/−^) to MPXV clade IIa (WR 7-61 and US-2003) and IIb (MA-2022) isolates. All four mouse models were susceptible to clade IIa infection, leading to severe disease marked by decreased body temperature, weight loss, and lethality. In contrast, clade IIb infection produced minimal to mild disease at similar doses in all four murine models.

The clade IIb isolate produced severe disease (40% lethality) at only the highest dose (8.0 log_10_ PFU) in the most susceptible immunocompromised mouse model, C57BL/6 *Ifnar*^−/−^/*Ifngr*^−/−^. This is the first demonstration of lethal disease with clade IIb in a murine model. In addition, these data demonstrate that clade IIa is ∼100- to 100,000-fold more virulent than clade IIb and provide three additional murine models for investigating MPXV infection and pathogenesis.

**IMPORTANCE:** Mpox is an emerging human disease caused by four distinct MPXV subclades (Ia, Ib, IIa, and IIb). Despite genetic similarities, the case fatality rate varies considerably between the subclades: Ia (∼11%), Ib and IIa (∼4%), and IIb (∼0.2%).

Since 2022, multiple mpox outbreaks have occurred due to previously unrecognized subclades, leading to the declaration of two public health emergencies by the World Health Organization. This unprecedented global spread, coupled with the variation in severity of human disease, underscores the importance of research into the pathogenesis of emerging MPXV subclades. However, a critical limitation is the lack of suitable small animal models. This study identifies three additional murine models susceptible to MPXV clade II infection and demonstrates significant virulence differences between clade IIa and IIb. These models will enable rapid characterization of previously unrecognized subclades and will facilitate countermeasure development.

## INTRODUCTION

Monkeypox virus (MPXV) is a member of the genus *Orthopoxvirus* in the family *Poxviridae*. The genome comprises a linear double-stranded DNA molecule ∼197 to 200 kilobase pairs (kbp) in length with covalently enclosed ends encoding ∼190 proteins (1–4). Mpox is a zoonotic disease endemic in parts of Central and West Africa (2, 5). The natural cycle required for maintenance of MPXV in nature is unknown, however, studies suggest that squirrels, giant pouched rats, African dormice, and non-human primates (NHP) can serve as sylvatic reservoirs (6–16). MPXV was first discovered in 1958 in Denmark, in a colony of cynomolgus macaques exhibiting a disease similar to smallpox (17). In the subsequent decade, the virus was isolated from multiple outbreaks in non-human primate (NHP) colonies in the USA and Europe following the importation of infected animals from Africa (18–23).

The first human case of mpox disease was reported in a young boy unvaccinated against smallpox in 1970 in Zaire (now the Democratic Republic of the Congo) (24). In the following five decades, sporadic outbreaks have been reported in West and Central Africa (2, 5). Human transmission of MPXV occurs through dermal or respiratory routes, and can cause severe disease characterized by fever, headache, fatigue, rash, lymphadenopathy, cutaneous lesions, myocarditis, encephalitis, and death (2).

MPXV is divided into two clades, clade I (Central Africa or Congo Basin) and clade II (West Africa), based on phylogenetics, geography, and the severity of human disease (25–28). Since the first reported human case of mpox disease, clade I isolates have accounted for a vast majority of the human cases. Sporadic outbreaks between 1970 and 2020 produced ∼30,000 confirmed or suspected cases with a case fatality rate of up to ∼11% (5). In contrast, only 200 cases were reported for clade II during the same period with a case fatality rate of ∼4% (5). For more than half a century, human mpox outbreaks have been confined to Africa. However, in 2022, the largest mpox outbreak occurred and rapidly spread throughout the globe (29, 30). The genetic sequencing of the 2022 mpox isolates revealed that the outbreak was caused by previously unrecognized subclade IIb. Phylogenetic analysis of the outbreak isolates demonstrated a close relationship to endemic isolates from Nigeria that have been circulating since 1971 and were responsible for the travel acquired cases exported to Europe and Asia in the 2017 outbreak (31–37). To date, the outbreak has produced ∼100,000 confirmed cases across 122 countries with 220 reported deaths and a case fatality rate of ∼0.2% (38, 39). In contrast to previous outbreaks, 96% of infections were reported in men who have sex with men (MSM) (38, 39). This was the first recorded instance of mpox disease through sexual transmission and highlights the potential of MPXV to cause global outbreaks.

Following the 2022 global outbreak, multiple outbreaks have occurred due to yet another previously unrecognized clade Ib (40–44). Consequently, MPXV now comprises two main clades with each further subdivided into subclades: Ia, Ib, IIa, and IIb. The human case fatality rates among the four subclades vary considerably, with rates of ∼11% (Ia), ∼4% (Ib and IIa), and ∼0.2% (IIa). The differences in virulence between MPXV clades in humans warrant further investigation, however, a critical impediment is the scarcity of small animal models. Comprehensive studies investigating the susceptibility of laboratory mouse strains to MPXV infection have been conducted previously (6, 40). Only four mouse strains including suckling white mice, SCID-BALB/c, C57BL/6 *stat1^-/-^*, and CAST/EiJ were shown to be susceptible to MPXV infection (6, 45–48). Of these models, C57BL/6 *stat1^-/-^*and CAST/EiJ have been utilized to investigate viral pathogenesis as well as countermeasure development (48–50). In addition, the majority of MPXV animal studies have utilized clade I isolates (6–16, 26, 45–60).

Consequently, few publications have compared the pathogenesis of clade IIa and IIb isolates (6, 45). In this study, we investigated the virulence differences of two MPXV clade IIa (WR 7-61 and US-2003) and one clade IIb (MA-2022) isolates in CAST/EiJ mice, as well as three immunocompromised mouse strains (C57BL/6 *Ifnar ^-/-^*, C57BL/6 *Ifngr ^-/-^*, and C57BL/6 *Ifnar* ^-/-^/*Ifngr ^-/-^*).

## RESULTS

### *In silico* and *in vitro* characterization of stock challenge of clade II isolates

To assess the evolutionary relatedness of MPXV, representative sequences from clades I and II were downloaded, aligned, and used to generate a maximum-likelihood phylogenetic tree (Fig. 1). As expected, two main clades were observed with two branches consisting of clade I Central African isolates and clade II West African isolates. Clade II further branched into clade IIa and IIb, with clade IIa comprising the oldest West African MPXV isolates WR 7-61 1962) and Sierra Leone (1970). In contrast, clade IIb is comprised of endemic isolates from Nigeria and travel acquired MPXV infections exported abroad. The MA-2022 isolate was placed within clade IIb, sister to isolates from Singapore and the United Kingdom.

**Figure 1.**
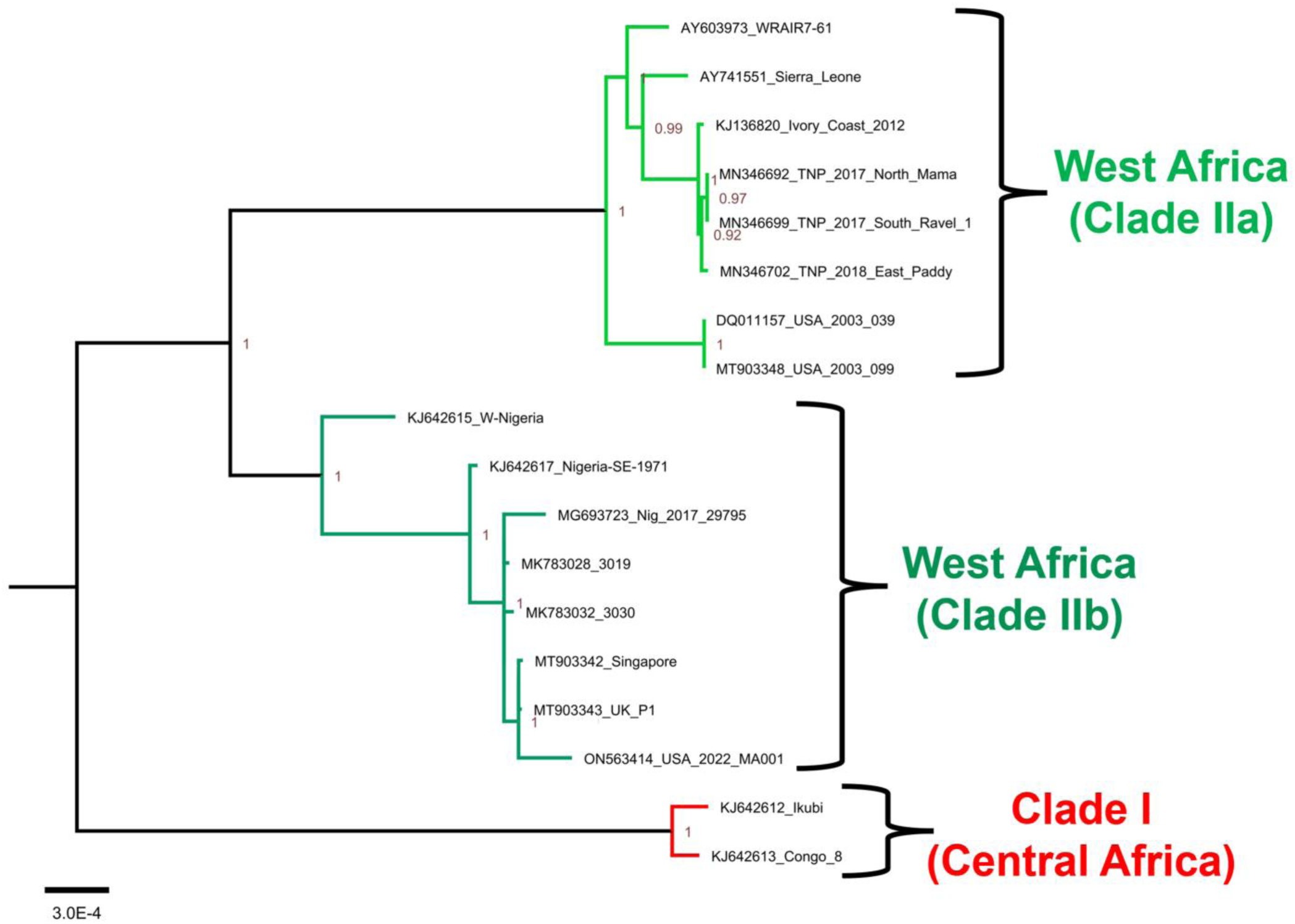
The phylogenetic relationship of MPVX isolates. Maximum likelihood tree based on the whole genome is shown with Central Africa (clade I – red), West Africa clade IIa (light green), and West Africa clade IIb (dark green).

Given the renewed interest in employing well-characterized representative isolates in validated animal models, we analyzed available sequence data from clade IIa and IIb MPXV isolates to document the variations in selected challenge stocks (61). Initially, we identified gene variations across MPXV clades and noted consistent clade-specific differences in gene length, truncation, and degradation (Table 1). Additionally, we documented gene polymorphisms that distinguished our selected challenge stocks from other clade members (Supp. Tables 1A and B). The phenotypic significance or potential correlations of these variations are beyond the scope of this work.

**Table 1.** Gene variations observed among MPXV clades.

| OPG | Observed variation | Name in VAC-Copenhagen | Clade Ia | Clade IIa | Clade IIb | Protein domain structure, localization, function | Clade IA aa Length | Clade IIa aa Length | Clade IIb aa Length | GB# GRI | CPXV-Bri | GRI aa Length |
| --- | --- | --- | --- | --- | --- | --- | --- | --- | --- | --- | --- | --- |
| **Genes missing in one or more clades** |  |  |  |  |  |  |  |  |  |  |  |  |
| OPG016 | Only present in Clade IIa | No homolog | NP | N3R | NP | MHC class I homolog, secreted, blocks NKG2D receptor, OMCP | 0 | 176 | 0 | CAD90591.1 | CPXV018 | 178 |
| OPG017 | Only present in Clade IIa | No homolog | NP | truncated N2 | NP | ANK and PRANC domains, binds to Cullin 2, inhibits NF-κB activation | 0 | 73 | 0 | CAD90592.1 | CPXV019 | 833 |
| OPG005 | Only present in Clade IIa | B22R/C16L | NP | N1R | NP | Bcl-2 domain, host range factor | 0 | 153 | 0 | CAD90589.1 | CPXV009 | 153 |
| OPG032 | Only present in Clade Ia | C3L | D14L | NP | NP | Complement control protein (CCP), prevents complement activation, secreted | 216 | 0 | 0 | CAA64102.1 | CPXV034 | 259 |
| OPG033 | Not present in MPXV | C2L | Degraded | NP | NP | BTB and Kelch domains, inhibits inflammation, reduces immunopathology | 0 | 0 | 0 | CAA64103.1 | CPXV035 | 512 |
| OPG033 | Only present in Clade Ia | C2L | D15L? | NP | NP |  | 105 | 0 | 0 |  |  |  |
| OPG033 | Only present in Clade Ia | C2L | D16L? | NP | NP |  | 77 | 0 | 0 |  |  |  |
| OPG033 | Only present in Clade Ia | C2L | D17L? | NP | NP |  | 98 | 0 | 0 |  |  |  |
| OPG033 | Only present in Clade Ia | C2L | D18L? | NP | NP |  | 107 | 0 | 0 |  |  |  |
| OPG195 | Only present in Clade Ia | B9R | B10R | Truncated | B10R | PIE, ER localized, retains MHC I in ER, prevents antigen presentation | 221 | 72 | 221 | CAD90734.1 | CPXV203 | 221 |
| OPG201 | Only present in Clade Ia | B16R | B14R | NP | NP | 3 immunoglobulin domains, IL-1 receptor homolog, blocks IL-1b, prevents fever in mice | 326 | 180 | 0 | CAD90740.1 | CPXV209 | 326 |
| OPG202 | Only present in Clade Ia | B17L | B15L | NP | NP | Distant homolog of A51 protein | 78 | 0 | 0 | CAD90741.1 | CPXV210 | 340 |
| **Genes varying in length between clades** |  |  |  |  |  |  |  |  |  |  |  |  |
| OPG029 | Polymorphism, STR repeat | C6L | D11L | D11L | D11L | Bcl-2 domain, inhibits type I IFN production and signaling, targets HDAC5 for degradation | 153 | 158 | 155 | CAA64099.1 | CPXV030 | 156 |
| OPG037 | Polymorphism, STR repeat | M1L | O1L | O1L | O1L | ANK, no PRANC domain, inhibits intrinsic apoptosis at the level of the apoptosome | 441 | 445 | 442 | CAA64107.1 | CPXV039 | 474 |
| OPG153 | Polymorphism, STR repeat | A26L | A28L | A28L | A28L | P4c precursor; A28L; major component of IMV surface tubules; p4c | 512-513 | 509-510 | 499-511 | CAD90694.1 | CPXV159 | 518 |
| OPG204 | Polymorphism, STR repeat | B19R | B16R | B16R | B16R | 3 immunoglobulin domains, IFN type-1 decoy receptor, secreted, attaches to GAGs | 350 | 354 | 356 | CAD90743.1 | CPXV212 | 351 |
| OPG208 | Polymorphism, STR repeat | K2L | B19R | B19R | B19R | Serpin 1,2,3; SPI-1; B19R; serine protease inhibitor-like, SPI-1; apoptosis inhibition | 389-392 | 524-542 | 449-497 | CAD90746.1 | CPXV217 | 375 |
| **Genes with variation in length in all MPXV (likely non-functional)** |  |  |  |  |  |  |  |  |  |  |  |  |
| OPG020 | Truncated in MPXV | C10L | Truncated | Truncated | Truncated | N-term prolyl hydroxylase fold, C-term IL-1 receptor antagonist, binds Ku | 83 | 67 | 37 | CAD90594.1 | CPXV022 | 331 |
| OPG027 | Truncated in MPXV | C7L | Truncated | Truncated | Truncated | Host range, inhibits type I IFN production and signaling | 64 | 105 | 72 | CAA64098.1 | CPXV029 | 150 |
| OPG041 | Truncated in MPXV | C3L | Truncated | Truncated | Truncated | Mimic of eIF2a, pseudo-substrate for PKR, PKR inhibitor, IFN resistance | 43 | 43 | 43 | CAA64111.1 | CPXV043 | 88 |

The plaque phenotype of all three isolates was investigated in Vero and BSC-40 cells (Fig. 1A). By 4 dpi, all three isolates produced plaques ∼1 to 2 mm and ∼2 to 3 mm in diameter on Vero and BSC-40 cells, respectively. In both cell lines, WR 7-61 and US-2003 isolates produced a predominantly uniform plaque phenotype, whereas MA-2022 isolate yielded a mixed plaque phenotype. The isolates were further analyzed by multiple-step growth kinetics in BSC-40 cells. All three isolates displayed similar replication kinetics with a ∼10,000-fold increase in titer by 24 hpi and achieved a peak titer of ∼8.0 log_10_ PFU/mL at 72 hpi (Fig. 2B).

**Figure 2.**
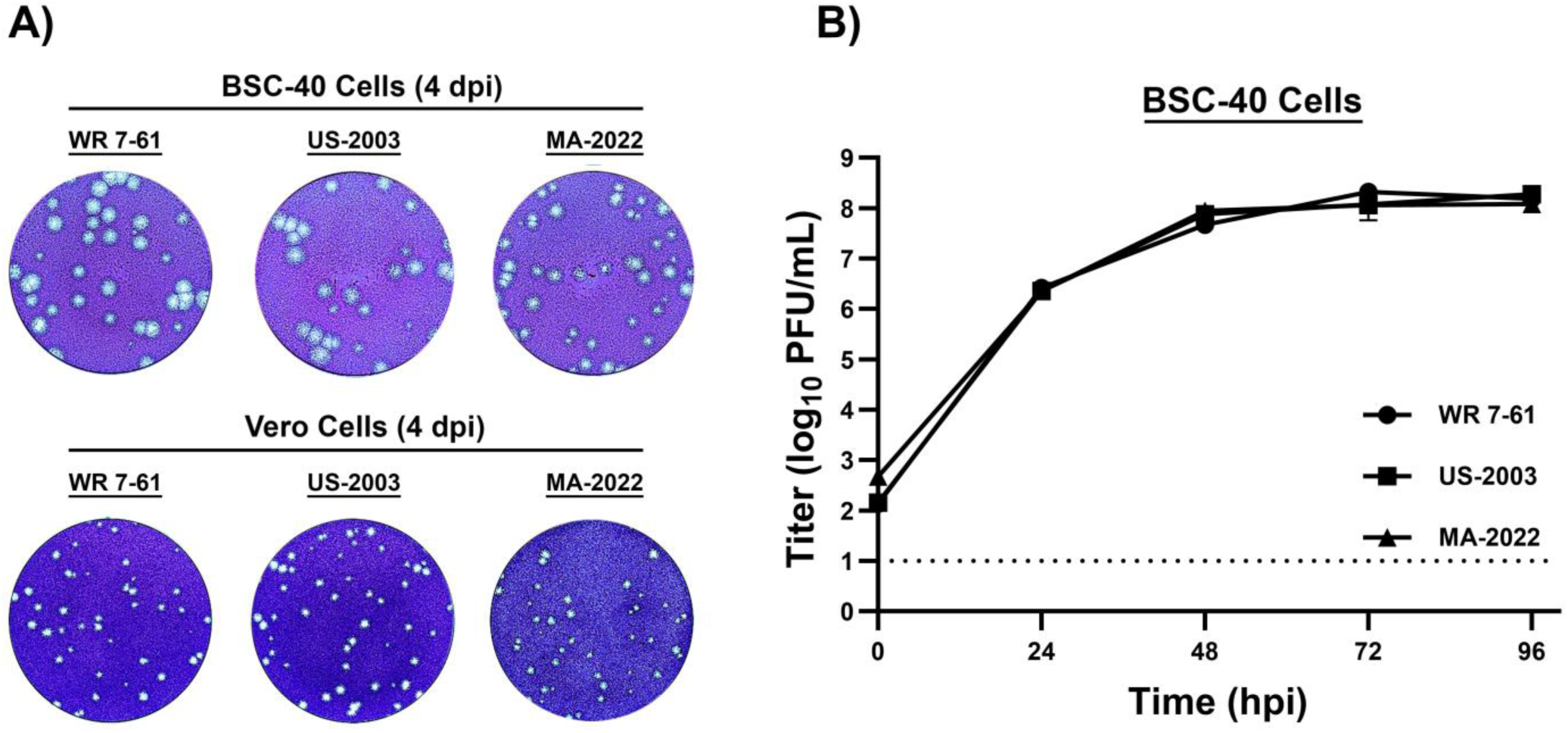
Plaque phenotype and multi-step growth kinetics of MPXV clade IIa isolates (WR 7-61 and US-2003) and IIb isolate (MA-2022). Plaque phenotype of MPXV isolates WR 7-61, US-2003, and MA-2022 was investigated in BSC-40 and Vero cells (A). Representative wells from 6-well plates at 4 dpi are shown. Multi-step growth kinetics of MPXV isolates in BSC-40 cells at an MOI of 0.01 (B). Samples were collected at 0-, 24-, 48-, 72-, and 96-hpi. Triplicate six-wells were sampled at each time-point followed by three cycles of freeze/thaw/sonication and titrated on BSC-40 cells. The limit of detection is shown for infectious virus (1.0 log_10_ PFU/mL) indicated with black dashed lines.

### Phenotypic difference between MPXV clade IIa (WR 7-61) and IIb (MA-2022) isolates in seven- to eight-week-old CAST/EiJ mice via the intranasal route

Cohorts of six to ten mice were infected with WR 7-61 (7.0, 5.0, 4.0 log_10_ PFU/mouse), MA-2022 (7.0 log_10_ PFU/mouse), or were mock-treated (Fig. 3A). Following infection, mice were monitored daily for disease score, body temperature, weight loss, and survival for 14 days. Severe disease was observed in mice infected with the WR 7-61 isolate in a dose-dependent manner. Mice infected at 7.0 log_10_ PFU exhibited increased disease score by 3 dpi with a peak average score of 2 at 6 dpi (Fig. 3B). Body temperature and weight declined beginning at 3 and 4 dpi, with peak average reduction of ∼-13 °F and ∼-20% at 6 dpi, respectively (Figs. 3C-D). All mice met euthanasia criteria by 7 dpi (Fig. 3E).

**Figure 3.**
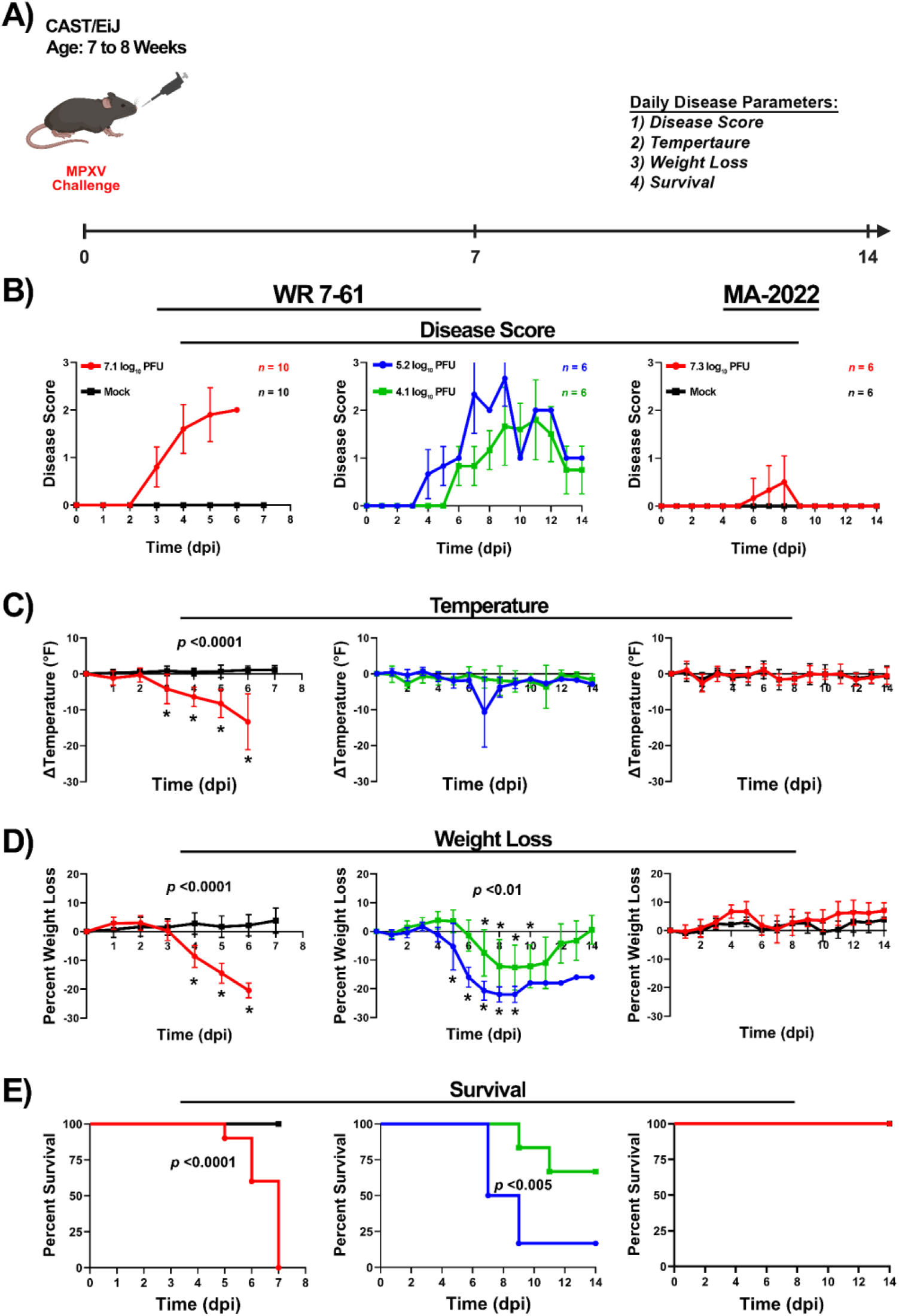
Infection of 7- to 8-week old CAST/EiJ mice with MPXV clade IIa (WR 7-61) and IIb (MA-2022) isolates via the intranasal (IN) route. Mice were infected with WR 7-61 at 7.0, 5.0, and 4.0 log_10_ PFU per mouse and with MA-2022 at 7.0 log_10_ PFU per mouse (A). Number of mice per group (n) ranged from 6 to 10 and is indicated for respective dose group (B). Following infection, mice were monitored for disease score (B), body temperature (C), weight loss (D), and survival (E). The target dose for each virus was verified by plaque assay, and values are shown (A). \**p*-values comparing MPXV isolates with the mock group are provided in each graph.

Mice infected with WR 7-61 at 5.0 and 4.0 log_10_ PFU/mouse exhibited increased disease scores between 4 and 6 dpi with peak average scores of ∼2.7 and ∼1.8 at 9 and 11 dpi, respectively (Fig. 3B). Alterations in body temperature were not observed in either dose group except at 7 dpi in the 5.0 log_10_ PFU group with mice exhibiting peak average temperature decline of ∼-11 °F (Fig. 3C). Body weight declined in both dose groups between 5 and 6 dpi with a peak average reduction of ∼-22% (5.0 log_10_ PFU) and ∼-13% (4.0 log_10_ PFU) (Fig. 3D). Eighty-three and thirty-three percent of the mice in 5.0 and 4.0 log_10_ PFU met euthanasia criteria, respectively (Fig. 3E).

In contrast to WR 7-61 infected mice, minimal disease was observed in mice infected with MA-2022. Disease scores increased between 6 and 8 dpi, with peak score of ∼0.5 (Fig. 3B). Minimal or no alterations were observed in body temperature or weight, and all mice survived the 14-day study (Figs. 3C-E).

### Phenotypic differences of MPXV clade IIa (WR 7-61) and IIb (MA-2022) isolates in four- to five-month-old CAST/EiJ mice via the intranasal route

Cohorts of five to six mice were infected with WR-7 61 (6.0 or 4.0 log_10_ PFU/mouse), MA-2022 (7.0 log_10_ PFU/mouse), or were mock-treated (Fig. 4A). WR 7-61 infected mice exhibited increased disease scores at 5 and 6 dpi, with peak average scores of ∼1.8 and ∼1.3 at 8 and 10 dpi (Fig. 4B). No alteration in body temperature was observed in the low dose group. However, in the higher dose group, mice exhibited a decrease in body temperature between 8 and 9 dpi, with a peak average decline of ∼-8 °F (Fig. 4C). Body weight loss was observed in both dose groups with peak average weight loss at 8 dpi of ∼-20% and ∼-10% in high- and low-dose groups, respectively (Fig. 4D). Fifty percent of mice infected with 6.0 log_10_ PFU met euthanasia criteria by 9 dpi, whereas all mice infected with 4.0 log_10_ PFU survived the 14-day study (Fig. 4E).

**Figure 4.**
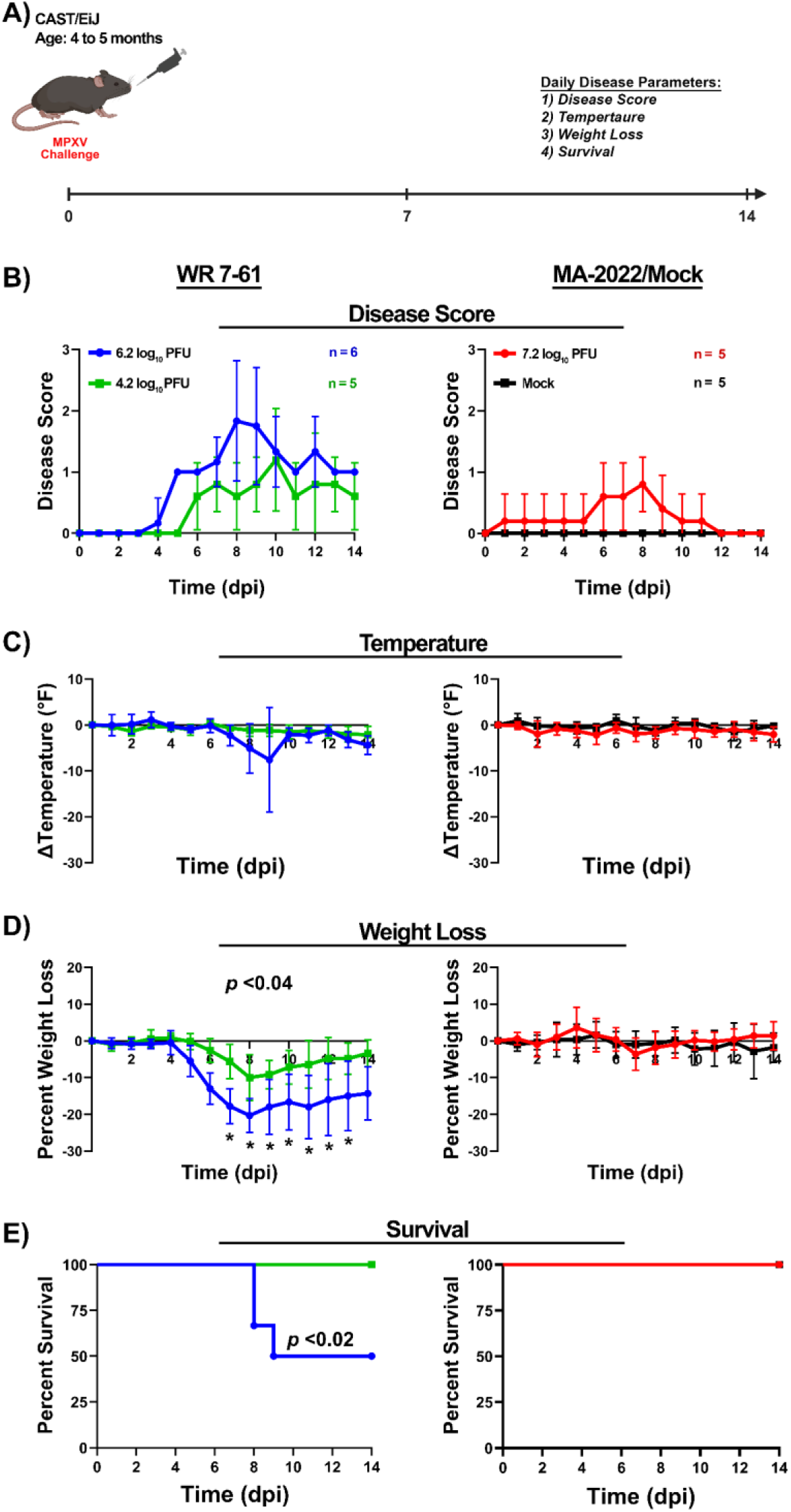
Infection of 4- to 5-month old CAST/EiJ mice with MPXV clade IIa (WR 7-61) and IIb (MA-2022) isolates via the intranasal (IN) route. Mice were infected with WR 7-61 at 6.0 and 4.0 log_10_ PFU per mouse and with MA-2022 at 7.0 log_10_ PFU per mouse (A). Number of mice per group (n) ranged from 5 to 6 and is indicated for respective dose group (B). Following infection, mice were monitored for disease score (B), body temperature (C), weight loss (D), and survival (E). The target dose for each virus was verified via the plaque assay and values are provided (A). \**p*-values comparing MPXV isolates with the mock group are provided in each graph.

In contrast to infection with WR 7-61, minimal disease was observed in mice infected with MA-2022. Disease scores increased between 1 and 11 dpi, with a peak average score of ∼0.8 (Fig. 4B). Minimal or no alterations were observed in body temperature and weight, and all mice survived the 14-day study (Figs. 4C-E).

### Phenotypic differences between MPXV clade IIa (WR 7-61, US-2003) and clade IIb (MA-2022) isolates in C57BL/6 *Ifnar*^−/−^ mice via the intranasal route

To investigate the susceptibility of C57BL/6 *Ifnar* ^−/−^ to MPXV infection, cohorts of 10 mice per group were infected via the intranasal route at 7.0, 6.0, or 5.0 log_10_ PFU/mouse (WR 7-61 and US-2003), 7.0 or 6.0 log_10_ PFU/mouse (MA-2022), or were mock-treated (Fig. 5A). As in previous studies, mice were monitored daily following infection for disease score, temperature, weight loss, and survival over 14 days. Disease scores rapidly increased within 3 dpi in mice infected with WR 7-61 and US-2003 at 7.0 and 6.0 log_10_ PFU/mouse dose groups and by 4 dpi all mice exhibited increased disease scores (Fig. 5B). The peak average disease scores were ∼1 to 2 between 5 to 9 dpi in all dose groups. Body temperature decreased rapidly within 2 dpi in all groups infected with WR 7-61 and US-2003, except US-2003 at 5.0 log_10_ PFU (Fig. 5C). The rate of temperature decline was dose-dependent with peak average decline of ∼-14 °F and ∼-10 °F in mice infected with WR 7-61 and US-2003 at 7.0 log_10_ PFU/mouse dose, respectively. In all other groups, peak average reduction ranged from ∼-3 °F to ∼-8 °F.

**Figure 5.**
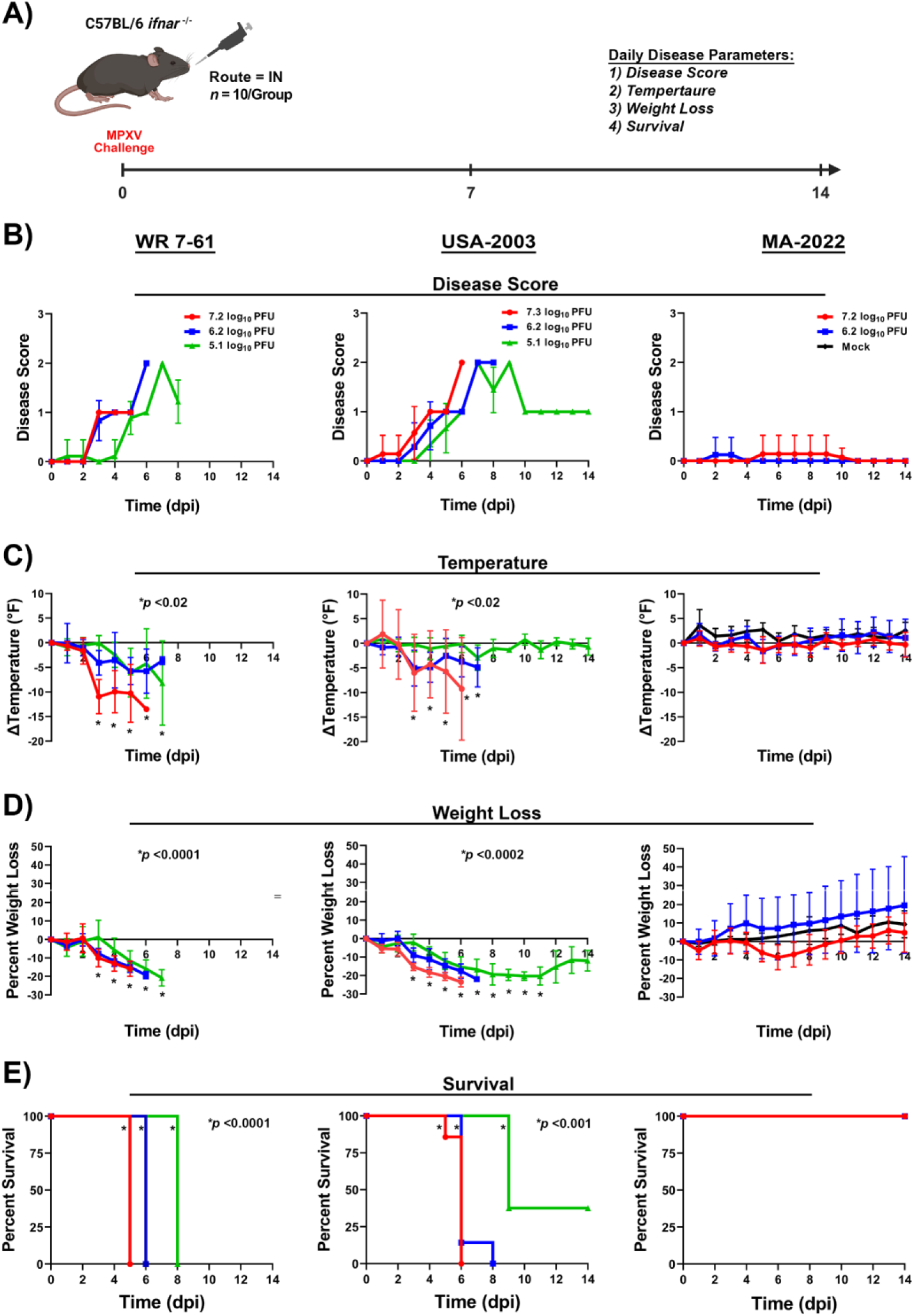
Infection of C57BL/6 *ifnar^-/-^*mice with MPXV clade IIa (WR 7-61 and US-2003) and IIb (MA-2022) isolates via the intranasal (IN) route. Mice were infected with WR 7-61 and US-2003 at 7.0, 6.0, and 5.0 log_10_ PFU per mouse and with MA-2022 at 7.0 and 6.0 log_10_ PFU per mouse (A). Following infection, mice were monitored for disease score (B), body temperature (C), weight loss (D), and survival (E). The target dose for each virus was verified by plaque assay, and values are shown (A). \**p*-values comparing MPXV isolates with the mock group are provided in each graph.

Mice infected with WR 7-61 and US-2003 isolates exhibited weight loss within 1 to 3 dpi (Fig. 5D). At 4 dpi, weight loss accelerated in all groups, with peak average weight loss of ∼-20% to ∼-24%. Only 40% of mice infected with US-2003 at 5.0 Log_10_ PFU/mouse regained weight, however, the weight gain plateaued by the end of the study. All mice infected with WR 7-61 at 7.0, 6.0, or 5.0 log_10_ PFU/mouse dose groups met euthanasia criteria by 5, 6, and 8 dpi, respectively (Fig. 5E). Similarly, all mice infected with US-2003 at 7.0 and 6.0 log_10_ PFU/mouse met euthanasia criteria at 5 to 6 dpi and 6 to 8 dpi, respectively. Partial lethality was observed at 5.0 log_10_ PFU/mouse group with 60% of the mice meeting euthanasia criteria by 9 dpi.

In contrast to WR 7-61 and US-2003 isolates, infection with MA-2022 was unable to produce severe disease in mice at any dose. Minimal or no changes were observed in disease score and body temperature (Figs. 5 B-C). Weight loss was only observed in the 7.0 log_10_ PFU/mouse group between 5 to 9 dpi, with peak average weight loss of ∼-9% (Fig. 5D). Mice in the lower dose groups gained weight throughout the 14-day study. All mice regardless of dose survived the infection with MA-2022 (Fig. 5E).

### Phenotypic differences between MPXV IIa (WR 7-61, US-2003) and IIb (MA-2022) isolates in C57BL/6 *Ifngr*^−/−^ mice via the intranasal route

All three MPXV isolates were next investigated in the C57BL/6 *Ifngr*^−/−^ background with a similar study design as previous studies (Fig. 6A). Mice infected with the two highest doses of WR 7-61 and US-2003 exhibited a rapid increase in disease score starting at 3 to 4 dpi with peak average score of ∼2 (Fig. 6B). The lowest dose groups displayed a 24-hr delay in onset of disease signs at 5 dpi. The peak average disease scores were ∼1.4 and ∼2.0 at 6 dpi and returning to baseline by 11 dpi. Mice infected with the two highest dose groups exhibited a decrease in body temperature starting at 3 dpi (Fig. 6C). Temperature decline peaked between 5 to 7 dpi with peak average decline of ∼-9 °F to -12 °F. In contrast, minimal or no decrease in body temperature was observed in mice infected with either isolate at the lowest dose.

**Figure 6.**
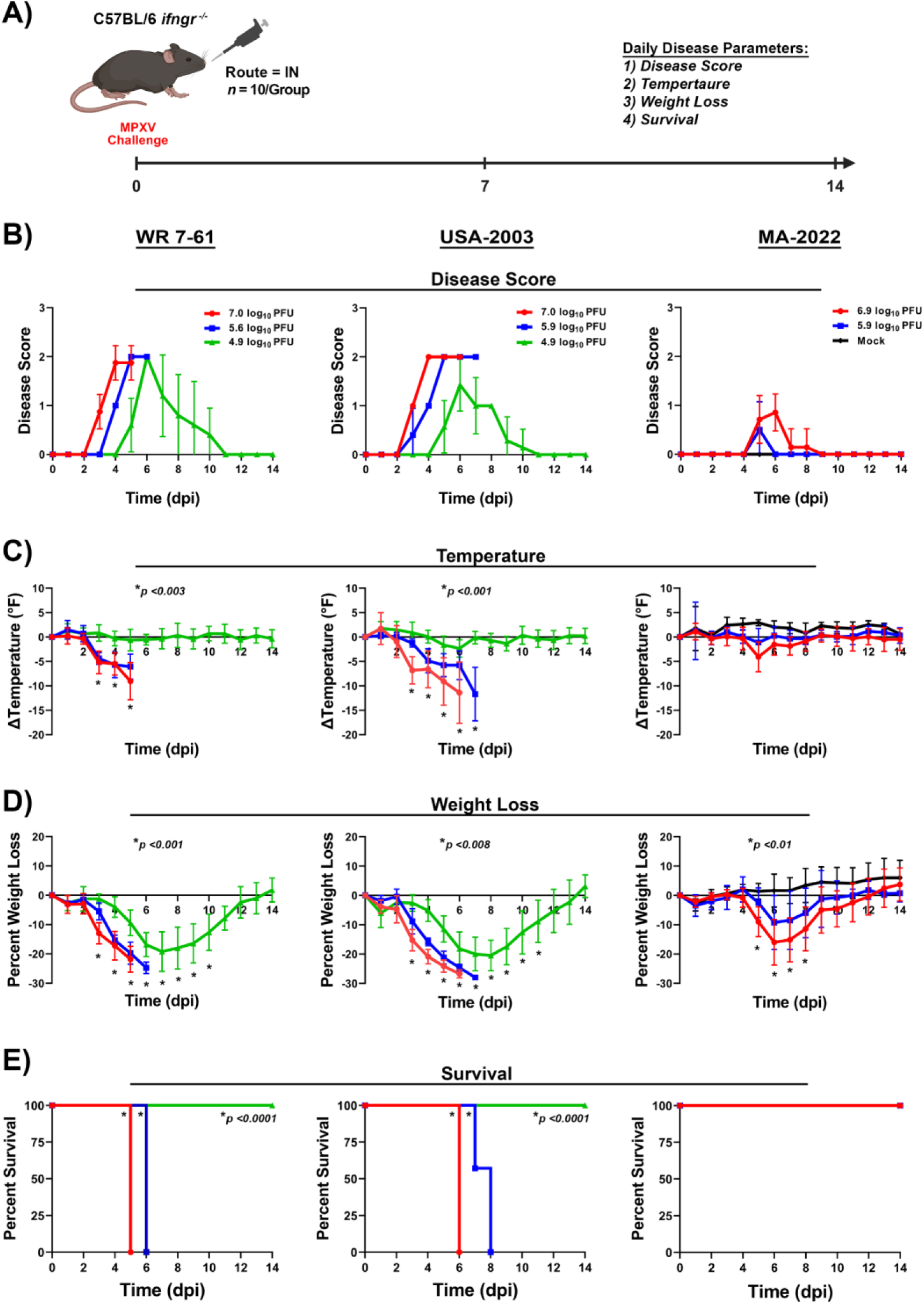
Infection of C57BL/6 *ifngr^-/-^*mice with MPXV clade IIa (WR 7-61 and US-2003) and IIb (MA-2022) isolates via the intranasal (IN) route. Mice were infected with WR 7-61 and US-2003 at 7.0, 6.0, and 5.0 log_10_ PFU per mouse and with MA-2022 at 7.0 and 6.0 log_10_ PFU per mouse (A). Following infection, mice were monitored for disease score (B), body temperature (C), weight loss (D), and survival (E). The target dose for each virus was verified by plaque assay, and values are shown (A). \**p*-values comparing MPXV isolates with the mock group are provided in each graph.

All mice infected with WR 7-61 or US-2003 exhibited weight loss, regardless of dose group (Fig. 6D). Mice exhibited weight loss starting at 1 dpi, with peak average weight loss of ∼-19% to -28% at 5 to 8 dpi. Mice in the lowest dose groups regained weight between 8 to 9 dpi and were at or above the initial weight by 14 dpi. All mice in two highest dose groups met euthanasia criteria between 5 and 8 dpi (Fig. 6E). In contrast, all mice infected with either isolate at the lowest dose survived the 14-day challenge study.

Mice infected with the MA-2022 isolate exhibited mild disease in both dose groups. Disease scores increased on 5 dpi, with peak average scores of ∼0.9 and 0.5, and returned to baseline by 9 dpi (high dose) and 6 dpi (low dose), respectively (Fig. 6B). Body temperature declined at the high dose between 5 to 8 dpi, with peak average decline of ∼-4 °F and by 9 dpi all mice returned to baseline (Fig. 6C). All mice in both dose groups exhibited weight loss in a dose-dependent manner beginning at 5 dpi, with peak average weight loss ranging from ∼-16% to -9% at 6 dpi (Fig. 6D). Mice began to regain weight by 7 dpi and returned to baseline by 12 (high group) and 10 dpi (low group), respectively. All mice infected with the MA-2022 isolate survived the 14-day study (Fig. 6E).

### Phenotypic differences between MPXV clade IIa (WR 7-61, US-2003) and IIb (MA-2022) isolates in C57BL/6 *Ifnar*^−/−^/ *Ifngr*^−/−^ mice via the intranasal route

The study design in C57BL/6 *Ifnar*^−/−^/*Ifngr*^−/−^ mice was similar to previous studies. Mice infected with WR 7-61 and US-2003 isolates received three doses (6.0, 5.0, and 4.0 log_10_ PFU/mouse), whereas MA-2022 infected mice received two doses (6.0 and 5.0 log_10_ PFU/mouse doses) (Fig. 7A). WR 7-61 and US-2003 infected mice at 6.0 log_10_ PFU/mouse exhibited severe disease, with rapid alterations in disease score, body temperature, and weight loss between 3 and 4 dpi (Figs. 7B-D). Disease score increased at 3 dpi, with a peak average score of ∼2 at 4 dpi. The peak decline in average body temperature ranged from ∼-7 °F and ∼-9 °F. Mice exhibited weight loss ranging from ∼-14% to -15%, and all mice met euthanasia criteria by 4 dpi.

**Figure 7.**
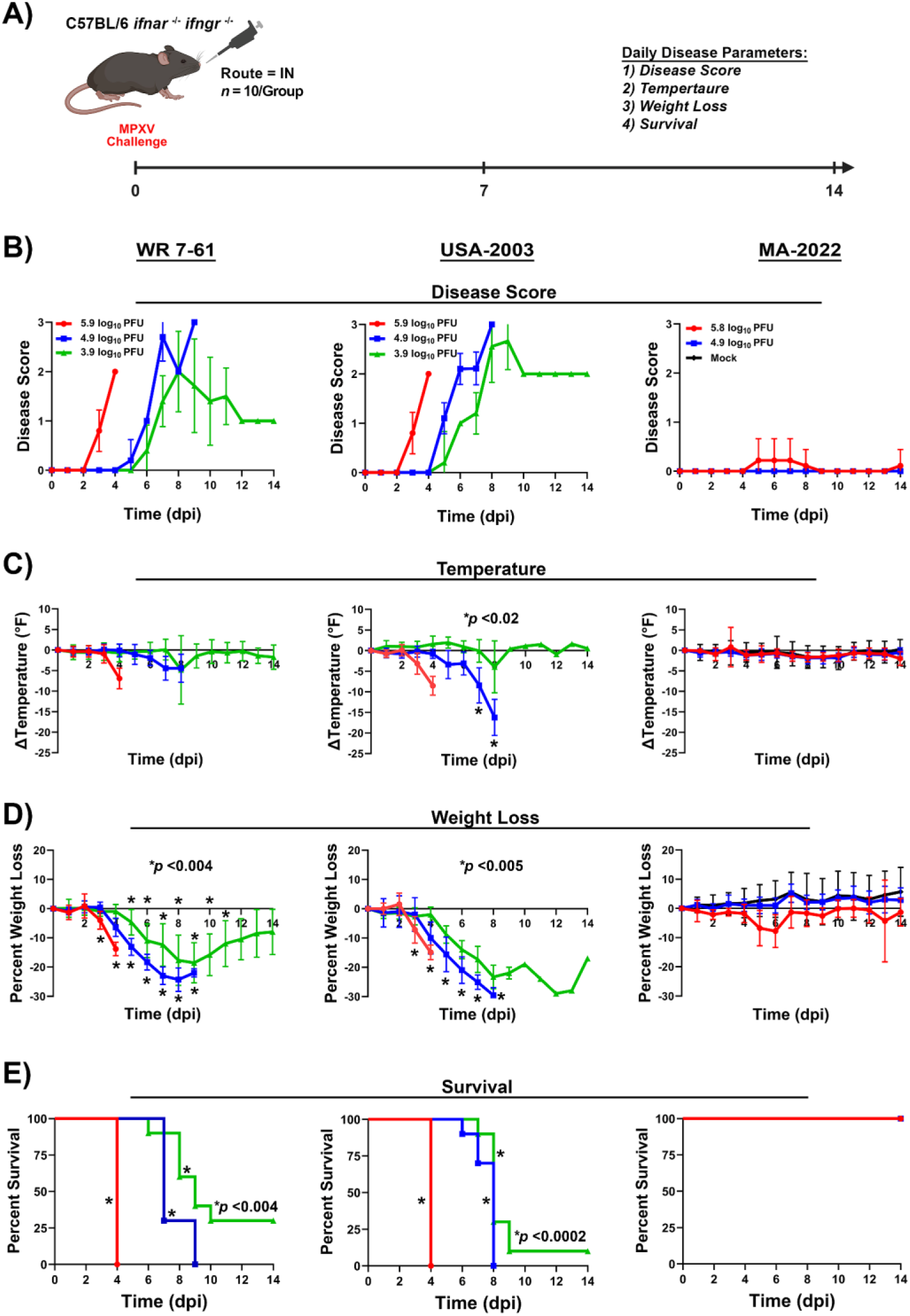
Infection of C57BL/6 *ifnar^-/-^/ifngr^-/-^* mice with MPXV clade IIa (WR 7-61 and US-2003) and IIb (MA-2022) isolates via the intranasal (IN) route. Mice were infected with WR 7-61 and US-2003 at 6.0, 5.0, and 4.0 log_10_ PFU per mouse and with MA-2022 at 6.0 and 5.0 log_10_ PFU per mouse (A). Following infection, mice were monitored for disease score (B), body temperature (C), weight loss (D), and survival (E). The target dose for each virus was verified by plaque assay, and values are shown (A). \**p*-values comparing MPXV isolates with the mock group are provided in each graph.

Mice infected with WR 7-61 and US-2003 at lower doses exhibited a delayed disease kinetics by 1 to 2 days. Disease scores increased between 5 and 6 dpi, with peak scores of ∼3 in the intermediate and ∼2 to 2.7 in the lowest dose groups (Fig. 7B). Body temperature began to decline between 6 and 8 dpi (Fig. 7C). In the intermediate-dose group, the temperature decline was more pronounced in US-2003 infected mice (∼−16 °F) than in WR 7-61 infected mice (∼−5 °F). The temperature decreased in the lowest dose groups at 8 dpi with average decline of ∼-5 °F, however, the temperature returned to baseline level within 24 hrs. Weight loss in intermediate and lowest dose groups exhibited similar kinetics, with the on-set of weight loss between 4 and 5 dpi (Fig. 7D). However, the magnitude of weight loss was greater in US-2003 infected mice than WR 7-61. At intermediate and lowest US-2003 dose groups, peak average weight loss was ∼-30%. In contrast, the intermediate and lowest WR 7-61 dose groups had weight loss of ∼-25% and ∼-19%, respectively. All mice infected with either isolate at the intermediate dose met euthanasia criteria between 6 and 9 dpi (Fig. 7E). At the lowest dose, 70% of WR 7-61–infected mice and 90% of US-2003 infected mice met euthanasia criteria between 6 and 10 dpi.

Similar to C57BL/6 *Ifnar*^−/−^ and *Ifngr*^−/−^ mouse studies, the MA-2022 isolate did not produce severe disease in C57BL/6 *Ifnar*^−/−^/*Ifngr*^−/−^ mice. Minimal or no changes were observed in disease score or body temperature (Figs. 7B-C). In the highest-dose group, modest weight loss of ∼-8% was observed between 4 and 5 dpi (Fig. 7D). The lowest dose group exhibited no weight loss and was indistinguishable from the mock infected group. All MA-2022 infected mice survived the 14-day challenge study (Fig. 7E).

A follow-up infection study in C57BL/6 *Ifnar*^−/−^/*Ifngr*^−/−^ mice was conducted at 3.0 and 2.0 log_10_ PFU (WR 7-61 and US-2003), and 8.0 log_10_ PFU (MA-2022) (Fig. 8A).

**Figure 8.**
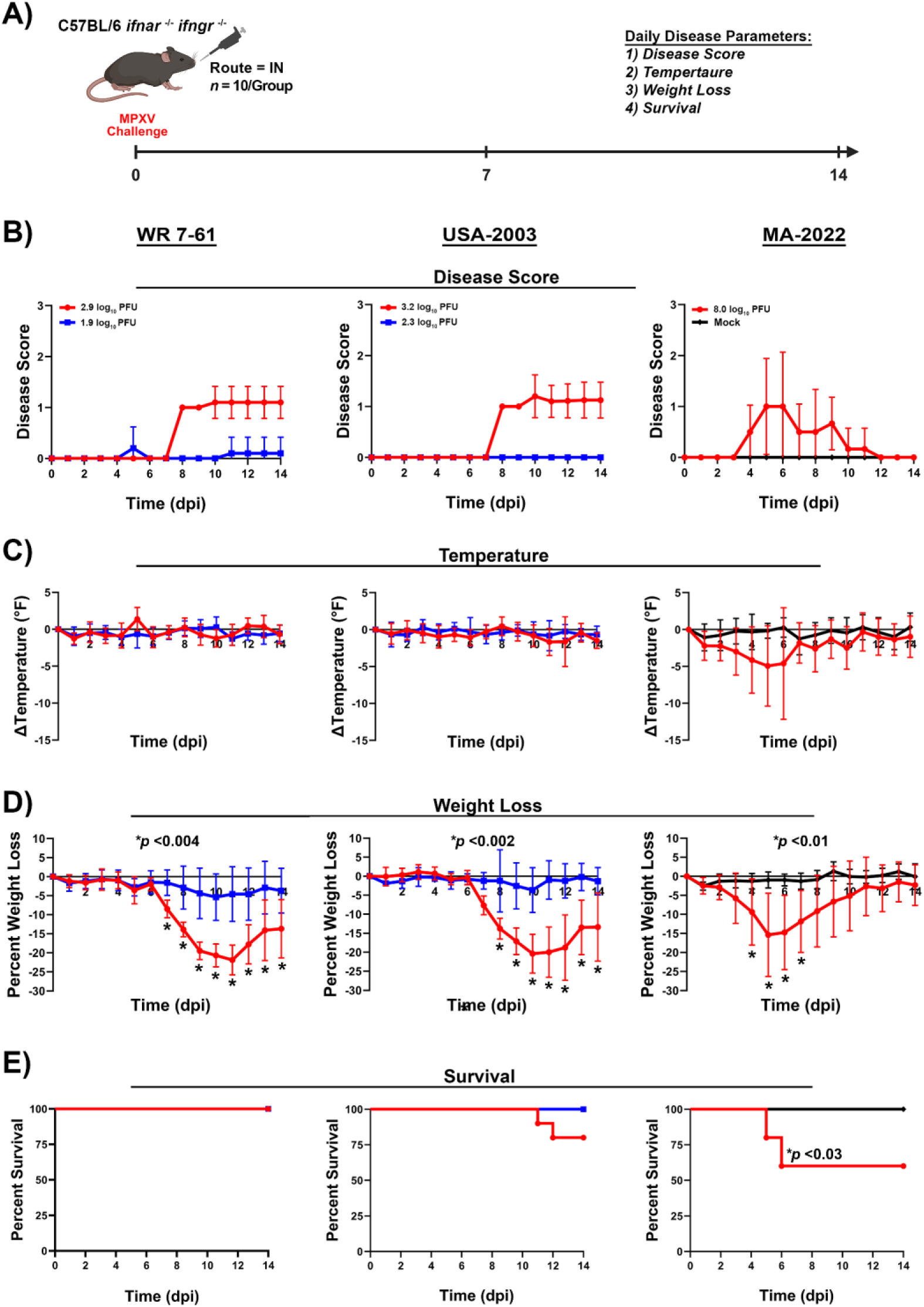
Infection of C57BL/6 *ifnar^-/-^/ifngr^-/-^* mice with MPXV clade IIa (WR 7-61 and US-2003) and IIb (MA-2022) isolates via the intranasal (IN) route. Mice were infected with WR 7-61 and US-2003 at 3.0 and 2.0 log_10_ PFU per mouse and with MA-2022 at 8.0 log_10_ PFU per mouse (A). Following infection, mice were monitored for disease score (B), body temperature (C), weight loss (D), and survival (E). The target dose for each virus was verified by plaque assay, and values are shown (A). \**p*-values comparing MPXV isolates with the mock group are provided in each graph.

Mice infected with WR 7-61 and US-2003 at 3.0 log_10_ PFU exhibited diseases scores from 7 to 14 dpi with peak scores of ∼1.2 (Fig. 8B). Only minimal alterations were observed in body temperature (Fig. 8C). Body weight declined at 7 dpi, with peak weight loss of ∼-21% between 10 to 11 dpi and did not return to baseline at 14 dpi (Fig. 8D). All mice infected with WR 7-61 survived the infection, whereas 20% of mice infected with US-2003 met the euthanasia criteria. Infection of mice with WR7-61 and US-2003 at a dose of 2.0 log_10_ PFU produced minimal disease. Minimal or no alterations in disease scores or body temperature were observed (Figs. 8B-C). A modest decrease in body weight was observed between 9 and 14 dpi, with peak decline by ∼-5% (Fig. 8D). All mice survived the 14-day infection study (Fig. 8E).

Mice infected with MA-2022 at 8.0 log_10_ PFU developed severe disease. Disease scores began to increase at 4 dpi and peaked at between 5 and 6 dpi, with average score of ∼1 between (Fig. 8B). Body temperature and weight started to decline at 1 and 3 dpi, with peak decline of ∼-5 °F and ∼-15% at 5 dpi (Figs. 8C-D). Forty percent of the mice met euthanasia criteria between 5 and 6 dpi (Fig. 8E).

### Susceptibility of C57BL/6 *Ifnar*^−/−^ mice to MPXV clade IIa (WR 7-61) and IIb (MA-2022) isolates via the intravaginal and intrarectal routes

C57BL/6 *Ifnar*^−/−^ mice were infected with WR 7-61 at 7.0, 6.0, 5.0, or 4.0 log_10_ PFU/mouse, MA-2022 at 7.0 log_10_ PFU/mouse, or mock treated via the intravaginal (IV) or intrarectal (IR) routes (Fig. 9A). Following infection with WR 7-61 at 7.0 log_10_ PFU/mouse via the IV route, mice exhibited redness and/or swelling at the site of challenge starting at 4 dpi. Eighty percent of mice exhibited vaginal redness and/or swelling with a peak average disease score of ∼1 at 8 dpi (Fig. 9B-C). The redness and swelling were largely resolved by 14 dpi. In contrast, no overt disease was observed in the lower dose groups. Body temperature varied by ∼-1 to -2 °F throughout the post-challenge period in all mice infected with WR 7-61, a pattern similar to that of the mock treated group (Fig. 9D).

**Figure 9.**
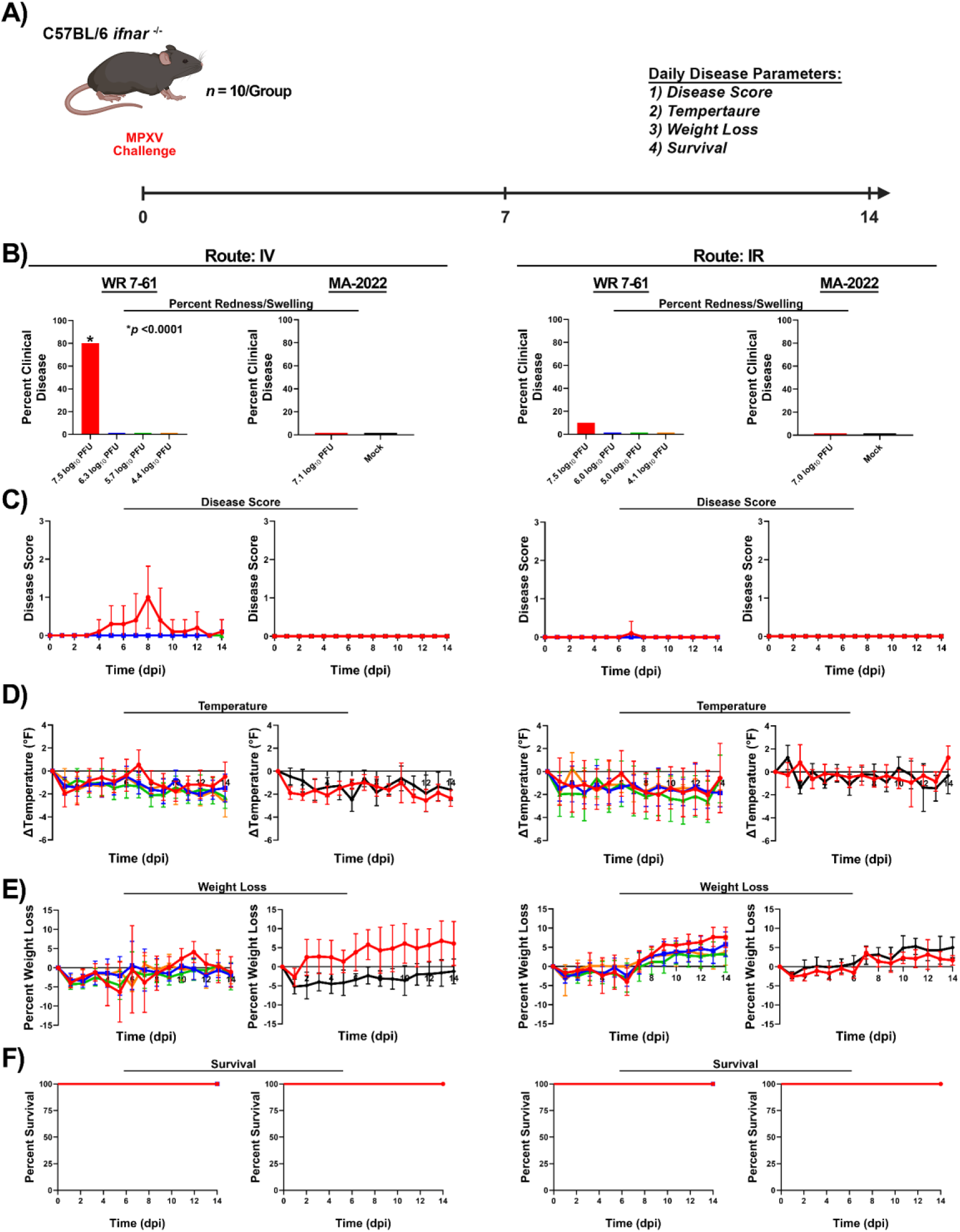
Infection of C57BL *ifnar^-/-^* mice with clade Ia (WR 7-61) and IIb (MA-2022) MPXV isolates via the intravaginal (IV) and intrarectal (IR) routes. Mice were infected with WR 7-61 at 7.0, 6.0, 5.0, and 4.0 log_10_ PFU per mouse and with MA-2022 at 7.0 log_10_ PFU per mouse (A). Following infection, mice were monitored for swelling and redness at challenge site (B), disease score (C), body temperature (D), weight loss (E), and survival (F). The target dose for each virus was verified by plaque assay, and values are shown (B). \**p*-values comparing MPXV isolates with the mock group are provided in each graph.

Mice exhibited weight loss in a dose dependent manner beginning at 2 dpi, with peak weight loss values of ∼-3% to -6% between 1 to 5 dpi (Fig. 9E). All mice, regardless of dose, survived the 14-day challenge study (Fig. 9F).

Infection of mice with WR 7-61 via the IR produced minimal or no redness/swelling at the site of infection, changes in disease score, or alterations in body temperature (Figs. 9B–D). However, mice did exhibit weight loss between 1 and 6 dpi with peak values ranging from ∼-2% to -4% (Fig. 9E). Similar to the IV challenge, all mice survived the 14-day challenge study (Fig. 9F).

Infection with MA-2022 via the IV or IR routes did not exhibit any overt disease or alteration in body temperature (Figs. 9B-D). Mice exhibited minimal weight loss and survived the 14-day study (Figs. 9E-F).

### Susceptibility of C57BL/6 *Ifnar*^−/−^/*Ifngr*^−/−^ mice to MPXV clade IIa (WR 7-61) and IIb (MA-2022) isolates via the intravaginal and intrarectal routes

The susceptibility via the IV and IR routes was investigated in C57BL/6 *Ifnar*^−/−^/*Ifngr*^−/−^ mice with WR 7-61 at 7.0, 6.0, 5.0, and 4.0 log_10_ PFU/mouse, MA-2022 at 7.0 log_10_ PFU/mouse, or mock treated (Fig. 10A). Twenty to eighty percent of mice infected with WR 7-61 via the IV route exhibited redness and/or swelling at the site of challenge (Fig. 10B). Mice infected with the highest dose exhibited disease scores throughout the 14-day challenge study, starting at 4 dpi with an average peak disease score of ∼1 (Fig. 10C). Infection at lower doses produced disease scores at 6 and 8 dpi, with average peak score of ∼0.5. Body temperature in all dose groups displayed minimal alterations and exhibited a similar pattern to that of the mock group (Fig. 10D). Body weight loss was observed in the highest dose group, with peak average weight loss of ∼-9% at 13 dpi (Fig 10E). Minimal weight loss was detected in all other dose groups. All mice regardless of dose survived the 14-day challenge study (Fig. 10F).

**Figure 10.**
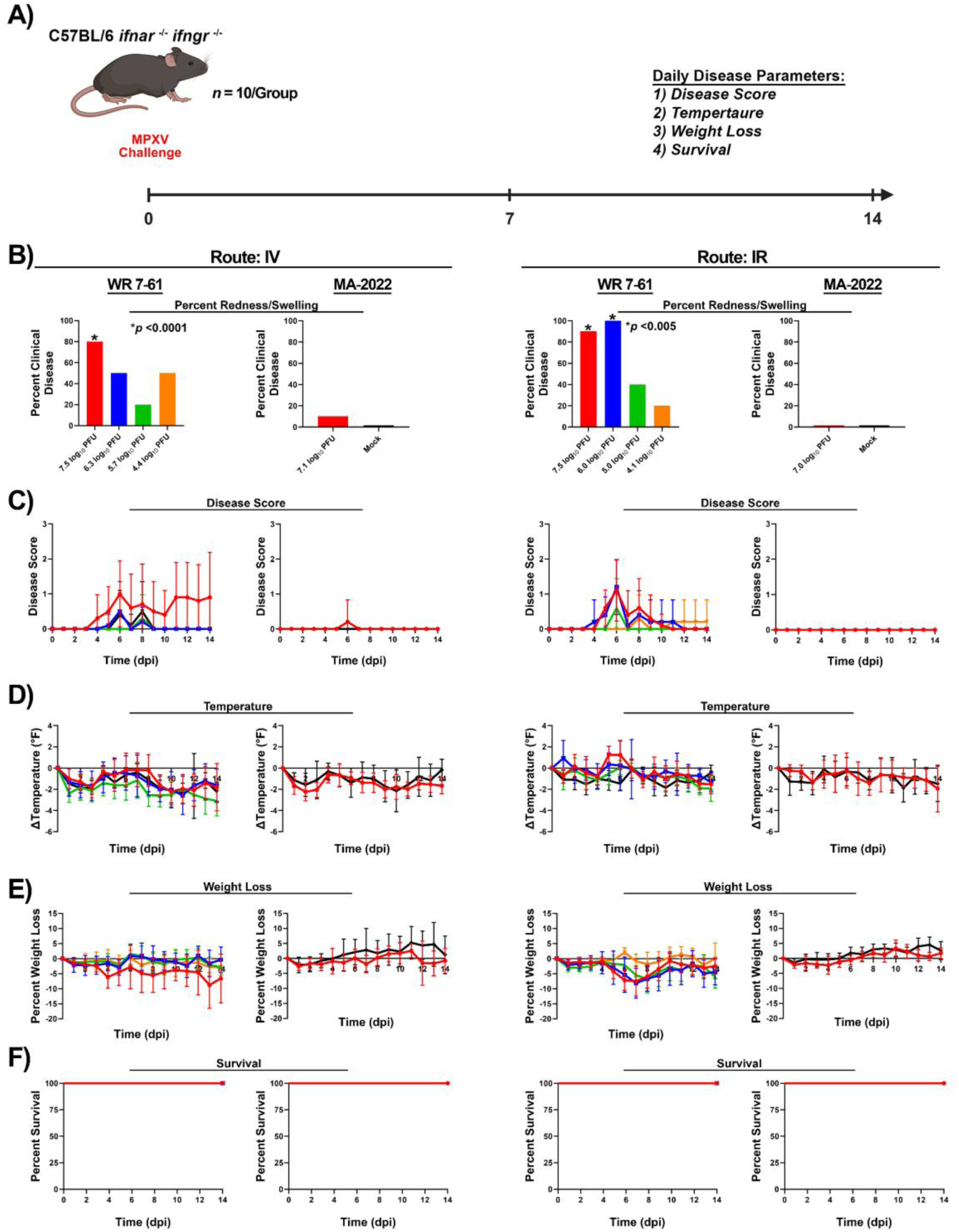
Infection of C57BL/6 *ifnar^-/-^/ifngr^-/-^* mice with clade Ia (WR 7-61) and IIb (MA-2022) MPXV isolates via the intravaginal (IV) and intrarectal (IR) routes. Mice were infected with WR 7-61 at 7.0, 6.0, 5.0, and 4.0 log_10_ PFU per mouse and with MA-2022 at 7.0 log_10_ PFU per mouse (A). Following infection, mice were monitored for swelling and redness at challenge site (B), disease score (C), body temperature (D), weight loss (E), and survival (F). The target dose for each virus was verified by plaque assay, and values are shown (B). \**p*-values comparing MPXV isolates with the mock group are provided in each graph.

Infection with the WR 7-61 isolate via the IR route produced redness and/or swelling in 20% to 100% of the mice in a dose-dependent manner (Fig. 10B). Mice exhibited disease scores between 4 and 14 dpi, with peak scores ranging from ∼0.3 to 1.2 (Fig. 10C). The pattern of variation in body temperature was similar to that of mock-infected mice across all dose groups (Fig. 10D). Body weight loss displayed a similar pattern in three of the highest dose groups (Fig. 10E). The mice exhibited weight loss starting at 5 dpi, with peak weight of ∼-7% to -8% between 7 and 8 dpi. Minimal weight loss was observed in the lowest dose group. Similar to the IV challenge study, all mice survived the 14-day challenge study (Fig. 10F).

In contrast to infection with WR 7-61, mice infected with MA-2022 via the IV or IR routes exhibited minimal or no disease, alteration in body temperature, minimal weight loss, and all mice survived the 14-day challenge study (Fig. 10B-F).

## DISCUSSION

Despite the threats that mpox poses to human health, few murine models are available for investigating the pathogenesis of emerging MPXV clades. Four mouse strains including suckling white mice, SCID-BALB/c, C57BL/6 *stat1^-/-^*, and CAST/EiJ have been shown to be susceptible to MPXV infection and exhibit lethal disease (6, 45–48). Of these models, the immune system of suckling mice and SCID-BALB/c is either not fully functional at the time of infection or is lacking. In contrast, both C57BL/6 *stat1^-/-^*and CAST/EiJ have deficiencies in the interferon response, which enables susceptibility to MPXV infection and disease. However, despite these deficiencies, both models can generate immune responses, and have been utilized in viral pathogenesis and countermeasure development studies (48–50).

Of the two murine models, only the CAST/EiJ has been used in previous comparative MPXV virulence studies (6, 45). The clade Ia Zaire-79-CB2 isolate (clonal or uncloned) was shown to be 2- to 10-fold more virulent than the clade IIa US-2003 isolate via the intranasal route (6, 45). The clade IIb isolate MA-2022 was unable to cause severe disease via either intranasal or intraperitoneal routes and was 100-fold less virulent than the US-2003 isolate (45). These studies demonstrated significant differences in the virulence of MPXV clades, with clade Ia exhibiting 1,000-fold greater virulent than clade IIb.

The present study investigated differences in the virulence of clade IIa (WR 7-61) and IIb (MA-2022) isolates in CAST/EiJ mice via the intranasal route. WR 7-61 isolate caused severe disease at 7.0, 5.0, and 4.0 log_10_ PFU in 7- to 8-week-old mice and at 6.0 log_10_ PFU in 4- to 5-month-old mice. In contrast, the MA-2022 isolate was unable to cause severe disease at 7.0 log_10_ PFU regardless of age. These data demonstrate that the clade IIb MA-2022 isolate is at least 1,000-fold less virulent than the clade IIa WR 7-61 isolate. In addition, these data corroborate previous clade II results and extend the findings in the CAST/EiJ model (40). Lastly, the susceptibility of 4- to 5-month-old CAST/EiJ mice to clade IIa infection highlights the utility of this model for evaluating long-term vaccine durability and efficacy.

Although the CAST/EiJ mouse model is susceptible to MPXV infection, it has several important limitations. First, commercial vendors have limited quantities, and large orders cannot be accommodated particularly during outbreaks or in projects with short timelines. Second, this mouse strain exhibits low pregnancy rates, small litter sizes, and high rates of maternal cannibalism. As a result of these traits, our attempt to establish an internal colony over approximately ∼1.5 years yielded poor results. Third, mice are highly energetic, and additional ABSL-2 and -3 personnel and safety procedures are required for breeding and experimental studies. For these reasons, only limited pilot studies with five to six animals per group were conducted, consequently warranting the investigation of additional mouse models.

Interferon alpha (IFN-α) and gamma (IFN-γ) play critical roles in the host response against poxvirus infection, and the disruption of either pathway dramatically increases the susceptibility of mice. Previous studies explored C57BL/6 *stat1^−/−^*mice that lack the STAT1 protein, which is essential for transmitting signals from type I and type II interferon receptors to the nucleus, resulting in a diminished antiviral immune response. Similarly, CAST/EiJ mice exhibit a deficient IFN-γ response, thus impairing an effective immune response against MPXV. In particular, a previous study investigated the susceptibility of CAST/EiJ mice to MPXV and found that the absence or reduced production of IFN-γ severely impaired the ability to control viral replication (49). As a result, MPXV spread unchecked, producing severe systemic infection and leading to lethality. The study also highlighted that CAST/EiJ mice failed to upregulate critical immune pathways dependent on IFN-γ, which is central to activating macrophages, promoting antigen presentation, and inducing antiviral states in infected cells. In a follow-up study delving further into the underlying mechanisms, the research demonstrated that these mice not only have an impaired IFN-γ response but also show insufficient innate immunity, characterized by low levels of Natural Killer (NK) cells following infection (50). NK cells are critical components of the early innate immune response, responsible for recognizing and eliminating virus-infected cells prior to the activation of the adaptive immune system. The mitigation of this defect via administration of NK cells or by the pre-stimulation of the interferon pathways led to improved outcomes. Pre-activation of interferon signaling reduced virus replication and disease severity while increasing survival rates. These studies highlight the importance of IFN in the host antiviral response against MPXV infection in mice.

Due to the critical role of IFN-α and IFN-γ, we investigated the susceptibility of three murine models, C57BL/6 *Ifnar*^−/−^, *Ifngr*^−/−^, and *Ifnar*^−/−^/*Ifngr*^−/−^, which lack functional type I and/or II interferon receptors. In addition to IFN deficiencies, these mouse strains are readily available through commercial vendors and are more amenable to the establishment of breeding colonies than CAST/EiJ mice. All three mouse strains were susceptible to clade IIa infection. The WR 7-61 and US-2003 isolates caused severe disease at doses ranging from 7.0 to 5.0 log_10_ PFU in C57BL/6 *Ifnar*^−/−^ and *Ifngr*^−/−^ mice. Not surprisingly, both isolates caused severe disease at 6.0 to 3.0 log_10_ PFU in the most immunocompromised murine model, C57BL/6 *Ifnar*^−/−^/*Ifngr*^−/−^ mice. In contrast to the clade IIa isolates, the clade IIb MA-2022 isolate did not cause severe disease at doses ranging from 7.0 to 5.0 log_10_ PFU. The MA-2022 isolate caused severe disease only at 8.0 log_10_ PFU dose in the C57BL/6 *Ifnar*^−/−^/*Ifngr*^−/−^. These data demonstrate that the C57BL/6 *Ifnar*^−/−^, *Ifngr*^−/−^, and *Ifnar*^−/−^/*Ifngr*^−/−^ mice are susceptible to MPXV clade II infection, and the clade IIb MA-2022 isolate from the 2022 outbreak is ∼100- to 100,000-fold less virulent than either clade IIa isolate (WR 7-61 or US-2003).

This study also investigated the susceptibility of C57BL/6 *Ifnar*^−/−^ and C57BL/6 *Ifnar*^−/−^/*Ifngr*^−/−^ mice to MPXV infection via the intravaginal and intrarectal routes. Infection with either WR 7-61 or MA-2022 was unable to produce severe disease in both mouse strain. However, mice infected with WR 7-61 exhibited weight loss and redness/swelling at the sites of infection, particularly via the intrarectal route. The redness/swelling was more pronounced in C57BL/6 *Ifnar*^−/−^/*Ifngr*^−/−^ mice at the infection site via both routes. In contrast, mice infected with MA-2022 did not exhibit any overt disease. These data further highlight the differences in virulence between clade IIa and IIb isolates.

IFN-α and/or IFN-γ receptor knockout mice have been previously investigated and were reported to be resistant to MPXV clade Ia infection (48). The data from the present study do not support the conclusions of the prior study. Several potential explanations may account for these conflicting results. First, the dose of MPXV clade Ia utilized in the previous study was too low (∼3.0 log_10_ PFU) to investigate susceptibility. Second, the receptor knockout approaches may not have completely eliminated interferon receptor function, and the residual activity enabled resistance to infection.

Third, the knockout mouse strains may have exhibited elevated basal levels of cytokines such as TNF-α, IL-1, and MIP-1α, which could have limited orthopoxvirus replication (62). However, these hypotheses require further investigation to reconcile the inconsistencies between the murine models.

To investigate the genetic basis of phenotypic differences among MPXV clades Ia, IIa, and IIb, we performed a comprehensive comparative genomic analysis (Table 1). Orthopoxviruses, including cowpox, horsepox, monkeypox, vaccinia, and variola comprise of large genomes ranging from ∼167 to 225 kbp dsDNA genomes encoding ∼170 to 214 genes (1, 63). The central ∼100 kbp core encodes genes responsible for virus replication and is consequently highly conserved. In contrast, the ∼30 to 40 kbp on either left or right arm of the genome are more divergent and encode host range restriction and host immune antagonism genes. The more virulent clade Ia encodes multiple host antagonism genes (D14L and B14R) that are absent or truncated in both clade IIa and IIb. D14L (C3L ortholog) encodes a complement binding protein that is a potent inhibitor of the complement-mediated antiviral response (64–67). In addition, the B14R gene encodes an IL-1 receptor homolog that inhibits host inflammatory response, is only present in clade Ia (68). Clade IIa encodes two additional virulence factors, N3R (OPG0165) and N1R (OPG005), which are absent in the less virulent clade IIb. N3R encodes a secreted MHC class I homolog that binds to and blocks the function of the NKG2D receptor, thereby reducing the activation of Natural Killer and T-cells (69). N1R encodes a protein with homology to the anti-apoptotic BCL-2 domain (70, 71). The presence or absence of these genes may contribute to the phenotypic differences between clades Ia, IIa, and IIb. In addition to these genes, comparative analyses have identified amino acid substitutions, insertions, and deletions, the significance of which is difficult to assess without comprehensive studies. Nonetheless, further experimental genetic studies are required to fully elucidate critical gene/s that underlie the phenotypic difference between MPXV clades.

There are several important limitations to our study. First, the virulence of clades Ia and Ib was not investigated in immunocompromised models. Clade I is more virulent than clade II in mice and would presumably exhibit greater virulence in the immunocompromised models. Second, whether immunocompromised model can be utilized in the context of mpox vaccine studies remains to be determined. Further studies are underway to address these limitations.

In summary, we investigated differences in the virulence of clade IIa and IIb isolates in CAST/EiJ, C57BL/6 *Ifnar*^−/−^, *Ifngr*^−/−^, and *Ifnar*^−/−^/*Ifngr*^−/−^ mice. In CAST/EiJ mice, the clade IIb MA-2022 isolate was up to 1,000-fold less virulent than clade IIa. Moreover, we demonstrated that three additional mouse models (C57BL/6 *Ifnar*^−/−^, *Ifngr*^−/−^, and *Ifnar*^−/−^/*Ifngr*^−/−^) are susceptible to MPXV clade II infection. Lastly, the MA-2022 isolate exhibited ∼100- to 100,000-fold lower virulence than clade IIa isolates in C57BL/6 *Ifnar*^−/−^, *Ifngr*^−/−^, and *Ifnar*^−/−^/*Ifngr*^−/−^ mice. Taken together, these data highlight differences in the virulence of clade II isolates and provide three additional murine models to investigate MPXV infection and pathogenesis.

## MATERIALS AND METHODS

### Phylogenetic analysis

Eighteen nonredundant genome sequences representing three main clades of MPXV were downloaded from Genbank and subsequently aligned with MAFFTv7 (FFT-NS-i strategy as iterative refinement method) (72). A Maximum Likelihood phylogenetic tree was reconstructed for the aligned dataset using PhyML under the GTR + GAMMA (4 CAT) substitution model (73).

### Cells

Vero (CRL-1586) and BSC-40 (CRL-2761) cells were obtained from American Tissue Culture Collection (ATCC, Manassas, VA). Cells were cultured in Dulbecco’s Modified Eagle Medium (DMEM) supplemented with 10% fetal bovine serum (FBS) and gentamicin (50 µg/mL) at 37°C and 5% CO_2_.

### Viruses and Virus Amplification

MPXV clade IIa (WR7-61 and US-2003) and IIb (MA-2022) isolates were obtained from BEI Resources (Manassas, VA). Virus stocks of each isolate were generated on BSC-40 cells. Briefly, cells were seeded overnight in flasks to achieve 70% confluence, growth media was removed, and monolayers were infected at a multiplicity of infection (MOI) of 0.01. Following an hour incubation at 37 °C and 5% CO_2_, 20 mL of growth media was added. Monolayers were harvested via cell scraping 72 hours post infection (hpi), samples were subjected to three cycles of freeze/thaw/sonication, and were stored at -80°C. For murine studies, virus stocks were sucrose purified. All virus stocks were titrated by plaque assay on BSC-40 cells.

### Virus Titration

BSC-40 cells were seeded overnight in 6-well plates to achieve 95-100% confluence. Each virus stock was serially diluted 10-fold in 1X DMEM. Monolayers were infected with 100 µL of each dilution and incubated at 37°C and 5% CO_2_ for one hour. Following incubation, monolayers were overlaid with 2 mL of 0.4% methylcellulose in MEM, GlutaMAX™ (Gibco) supplemented with 5% FBS and gentamicin (50 µg/mL). Following 72 hours at 37°C and 5% CO_2_, plates were fixed in 10% formalin overnight. Plates were stained with 2% crystal violet in 70% ethanol for 5-10 minutes. The stain was removed with water, dried, and plaques were counted. For plaque phenotype comparison, an identical protocol was performed with Vero and BSC-40 cells, and plaques were imaged 96 hpi.

### Multi-Step Growth Kinetics

BSC-40 cells were seeded overnight in 6-well plates to achieve 50% confluence. Growth media was removed, and triplicate monolayers were infected at an MOI of 0.01 with MPXV isolates. Following an hour incubation at 37°C and 5% CO_2_, monolayers were washed 3 times with 10 mL PBS, and 2 mL of growth media was added. Monolayers were harvested via cell scraping at 0-, 24-, 48-, 72-, and 96-hpi. Samples were subjected to three cycles of freeze/thaw/sonication and were stored at -80°C.

### Ethics statement

Research was conducted under an Institutional Animal Care and Use Committee (IACUC) approved protocol and in compliance with the Animal Welfare Act, PHS Policy, and other Federal statutes and regulations relating to animals and experiments involving animals. The facility where this research was conducted is USDA registered and adheres to principles stated in the Guide for the Care and Use of Laboratory Animals, National Research Council, 2011 [16].

### Mouse Studies

Breeding pairs of CAST/EiJ, C57BL/6 *ifnar ^-/-^*, *ifngr ^-/-^*, *ifnar ^-/-^*/*ifngr ^-/-^*were purchased from Jackson Laboratories to generate mouse colonies. Fourteen-day studies were performed to investigate virulence differences between the clade IIa (WR 7-61 and US-2003) and IIb (MA-2022) isolates. For CAST/EiJ studies, 7- to 8-week and 4- to 5-month-old mice were utilized with 5 to 10 mice per group. Mice were infected with WR 7-61 (7.0, 6.0, 5.0, and 4.0 log_10_ PFU) and MA-2022 (7.0 log_10_ PFU) via the intranasal route. Mice were infected with 30 µL (15 µL/nare) of virus or 1X PBS. For C57BL/6 *ifnar ^-/-^*, *ifngr ^-/-^*, *ifnar ^-/-^*/*ifngr ^-/-^*mouse studies, cohorts of ten (5 male and 5 female) six- to eight-week-old mice were infected with MPXV isolates via the intranasal route. Mice were infected with 30 µL (15 µL/nare) of virus or 1X PBS. C57BL/6 *ifnar ^-/-^*and *ifngr ^-/-^* mice were infected with three different doses, 7.0, 6.0, and 5.0 log_10_ PFU/mouse with WR 7-61 and US-2003 isolates. Both stains were also infected with MA-2022 at 7.0 and 6.0 log_10_ PFU/mouse. C57BL/6 *ifnar ^-/-^*/*ifngr ^-/-^* mice were infected with 6.0, 5.0, 4.0, 3.0, and 2.0 log_10_ PFU/mouse of WR 7-61 or US-2003 isolates. Mice were also infected with MA-2022 at 8.0, 6.0, and 5.0 log_10_ PFU/mouse. Following infection, all mice were monitored daily for disease score, body temperature, weight loss, and survival. The disease score criteria: 0 = normal, 1 = rough coat and/or hunched posture, 20% body weight loss from baseline, 2 = mild lethargy and/or mild dyspnea, 25% body weight loss from baseline, and 3 = moderate lethargy and/or moderate dyspnea and/or 30% body weight loss from baseline. All animal dosing was verified via plaque assay and are shown in respective figures.

### Statistical Analysis

Statistical significance in temperature, weight loss, and survival was evaluated using nonparametric one-way analysis of variance (ANOVA) with Tukey’s multiple comparison tests, non-parametric Mann-Whitney *t-test*, or unpaired multiple *t-tests* using a false discovery rate approach. A comparison of survival curves was performed using the Log-rank (Mantel-Cox) test, the Logrank test for trend, and Gehan-Breslow-Wilcoxon tests. The representative *p-values* for each analysis are provided in the figures. All the tests were performed using GraphPad PRISM software (version 10.0.1). Statistically significant *p*-values <0.05 are shown in the graphs.

---

**Supp. Table 1a.**
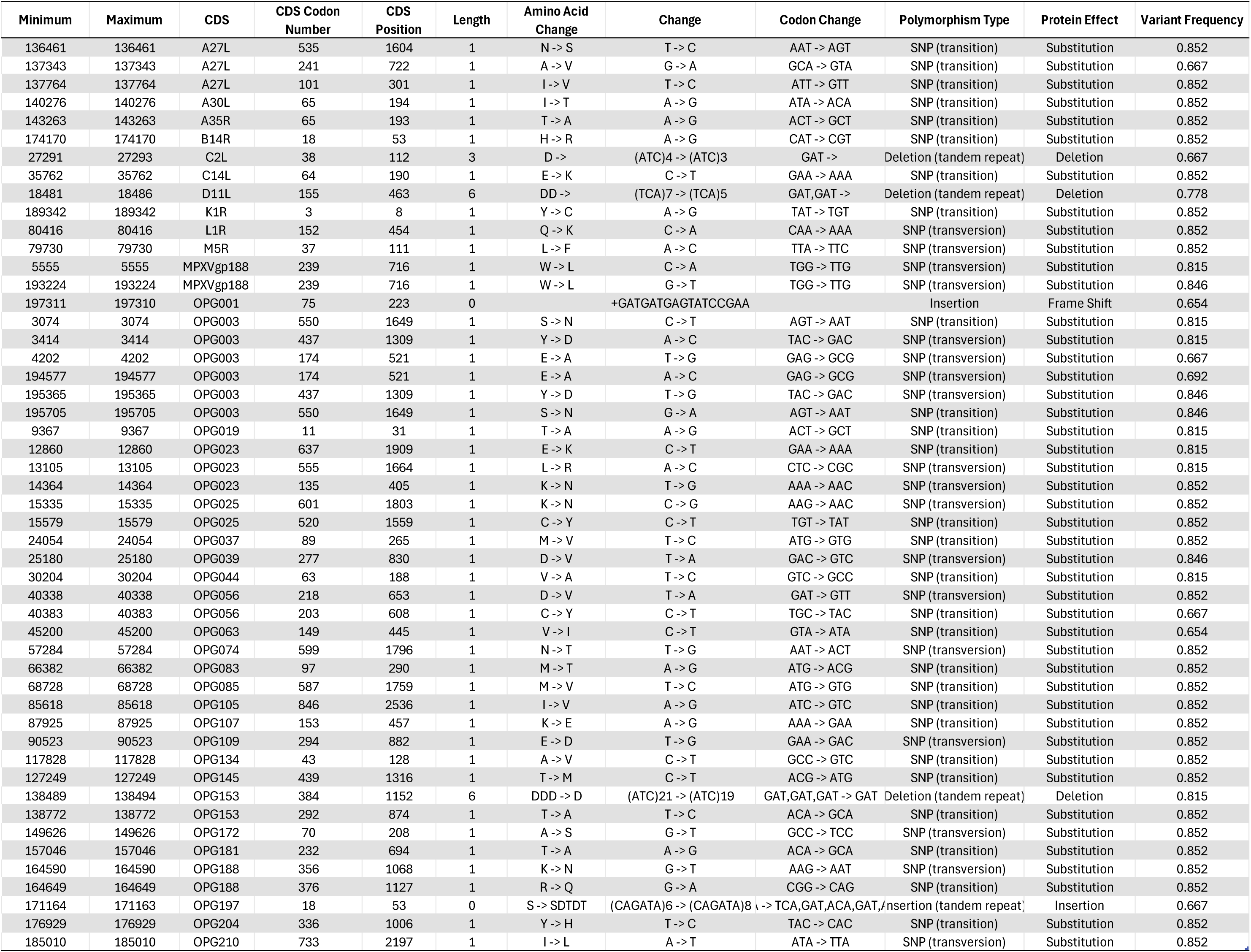
. Genetic polymorphisms between MPXV clade IIa Isolates (US-2003 and WR 7-61).

**Supp. Table 1b.**
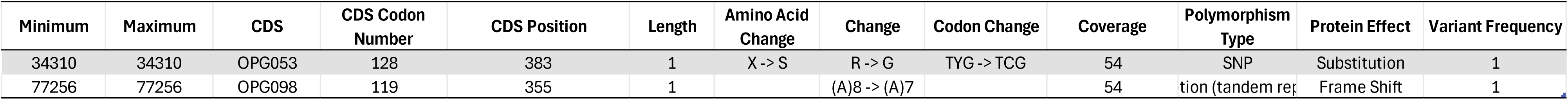
. Genetic polymorphisms between MPXV clade IIb isolates.

## References

1. Moss B, Smith GL. 2021. Poxviridae: The Viruses and Their Replication, vol 2. Lippincott Williams & Wilkins (LWW).

2. Panayampalli S. Satheshkumar, Damon IK. 2021. Poxviruses, vol 2. Lippincott Williams & Wilkins (LWW).

3. Shchelkunov SN, Totmenin AV, Babkin IV, Safronov PF, Ryazankina OI, Petrov NA, Gutorov VV, Uvarova EA, Mikheev MV, Sisler JR, Esposito JJ, Jahrling PB, Moss B, Sandakhchiev LS. 2001. Human monkeypox and smallpox viruses: genomic comparison. FEBS Lett 509:66–70.

4. Shchelkunov SN, Totmenin AV, Safronov PF, Mikheev MV, Gutorov VV, Ryazankina OI, Petrov NA, Babkin IV, Uvarova EA, Sandakhchiev LS, Sisler JR, Esposito JJ, Damon IK, Jahrling PB, Moss B. 2002. Analysis of the monkeypox virus genome. Virology 297:172–94.

5. Bunge EM, Hoet B, Chen L, Lienert F, Weidenthaler H, Baer LR, Steffen R. 2022. The changing epidemiology of human monkeypox-A potential threat? A systematic review. PLoS Negl Trop Dis 16:e0010141.

6. Americo JL, Moss B, Earl PL. 2010. Identification of wild-derived inbred mouse strains highly susceptible to monkeypox virus infection for use as small animal models. J Virol 84:8172–80.

7. Khodakevich L, Jezek Z, Kinzanzka K. 1986. Isolation of monkeypox virus from wild squirrel infected in nature. Lancet 1:98–9.

8. Hutson CL, Nakazawa YJ, Self J, Olson VA, Regnery RL, Braden Z, Weiss S, Malekani J, Jackson E, Tate M, Karem KL, Rocke TE, Osorio JE, Damon IK, Carroll DS. 2015. Laboratory Investigations of African Pouched Rats (Cricetomys gambianus) as a Potential Reservoir Host Species for Monkeypox Virus. PLoS Negl Trop Dis 9:e0004013.

9. Lourie B, Nakano JH, Kemp GE, Setzer HW. 1975. Isolation of poxvirus from an African Rodent. J Infect Dis 132:677–81.

10. Hutson CL, Lee KN, Abel J, Carroll DS, Montgomery JM, Olson VA, Li Y, Davidson W, Hughes C, Dillon M, Spurlock P, Kazmierczak JJ, Austin C, Miser L, Sorhage FE, Howell J, Davis JP, Reynolds MG, Braden Z, Karem KL, Damon IK, Regnery RL. 2007. Monkeypox zoonotic associations: insights from laboratory evaluation of animals associated with the multi-state US outbreak. Am J Trop Med Hyg 76:757–68.

11. Tesh RB, Watts DM, Sbrana E, Siirin M, Popov VL, Xiao SY. 2004. Experimental infection of ground squirrels (Spermophilus tridecemlineatus) with monkeypox virus. Emerg Infect Dis 10:1563–7.

12. Falendysz EA, Lopera JG, Doty JB, Nakazawa Y, Crill C, Lorenzsonn F, Kalemba LN, Ronderos MD, Mejia A, Malekani JM, Karem K, Carroll DS, Osorio JE, Rocke TE. 2017. Characterization of Monkeypox virus infection in African rope squirrels (Funisciurus sp.). PLoS Negl Trop Dis 11:e0005809.

13. Sbrana E, Xiao SY, Newman PC, Tesh RB. 2007. Comparative pathology of North American and central African strains of monkeypox virus in a ground squirrel model of the disease. Am J Trop Med Hyg 76:155–64.

14. Sergeev AA, Kabanov AS, Bulychev LE, Sergeev AA, Pyankov OV, Bodnev SA, Galahova DO, Zamedyanskaya AS, Titova KA, Glotova TI, Taranov OS, Omigov VV, Shishkina LN, Agafonov AP, Sergeev AN. 2017. Using the Ground Squirrel (Marmota bobak) as an Animal Model to Assess Monkeypox Drug Efficacy. Transbound Emerg Dis 64:226–236.

15. Schultz DA, Sagartz JE, Huso DL, Buller RM. 2009. Experimental infection of an African dormouse (Graphiurus kelleni) with monkeypox virus. Virology 383:86–92.

16. Radonic A, Metzger S, Dabrowski PW, Couacy-Hymann E, Schuenadel L, Kurth A, Matz-Rensing K, Boesch C, Leendertz FH, Nitsche A. 2014. Fatal monkeypox in wild-living sooty mangabey, Cote d’Ivoire, 2012. Emerg Infect Dis 20:1009-11.

17. Magnus Pv, Andersen EK, Petersen KB, Birch-Andersen A. 1959. A pox-like disease in cynomolgus monkeys. Acta Pathologica Microbiologica Scandinavica 46:156–176.

18. McConnell SJ, Herman YF, Mattson DE, Erickson L. 1962. Monkey pox disease in irradiated cynomologous monkeys. Nature 195:1128–1129.

19. Peters J. 1966. An epizootic of monkey pox at Rotterdam Zoo. International Zoo Yearbook 6:274–275.

20. Prier J, Sauer RM. 1960. Apox disease of monkeys. Annals of the New York Academy of Sciences 85:951–959.

21. Prier JE, Sauer RM. 1960. A pox disease of monkeys. Ann N Y Acad Sci 85:951–9.

22. Prier JE, Sauer RM, Malsberger RG, Sillaman JM. 1960. Studies on a pox disease of monkeys. II. Isolation of the etiologic agent. Am J Vet Res 21:381–4.

23. Sauer RM, Prier JE, Buchanan RS, Creamer AA, Fegley HC. 1960. Studies on a pox disease of monkeys. I. Pathology. Am J Vet Res 21:377–80.

24. Ladnyj ID, Ziegler P, Kima E. 1972. A human infection caused by monkeypox virus in Basankusu Territory, Democratic Republic of the Congo. Bull World Health Organ 46:593–7.

25. Likos AM, Sammons SA, Olson VA, Frace AM, Li Y, Olsen-Rasmussen M, Davidson W, Galloway R, Khristova ML, Reynolds MG, Zhao H, Carroll DS, Curns A, Formenty P, Esposito JJ, Regnery RL, Damon IK. 2005. A tale of two clades: monkeypox viruses. J Gen Virol 86:2661–2672.

26. Chen N, Li G, Liszewski MK, Atkinson JP, Jahrling PB, Feng Z, Schriewer J, Buck C, Wang C, Lefkowitz EJ, Esposito JJ, Harms T, Damon IK, Roper RL, Upton C, Buller RM. 2005. Virulence differences between monkeypox virus isolates from West Africa and the Congo basin. Virology 340:46–63.

27. Isidro J, Borges V, Pinto M, Sobral D, Santos JD, Nunes A, Mixao V, Ferreira R, Santos D, Duarte S, Vieira L, Borrego MJ, Nuncio S, de Carvalho IL, Pelerito A, Cordeiro R, Gomes JP. 2022. Addendum: Phylogenomic characterization and signs of microevolution in the 2022 multi-country outbreak of monkeypox virus. Nat Med 28:2220–2221.

28. Happi C, Adetifa I, Mbala P, Njouom R, Nakoune E, Happi A, Ndodo N, Ayansola O, Mboowa G, Bedford T, Neher RA, Roemer C, Hodcroft E, Tegally H, O’Toole A, Rambaut A, Pybus O, Kraemer MUG, Wilkinson E, Isidro J, Borges V, Pinto M, Gomes JP, Freitas L, Resende PC, Lee RTC, Maurer-Stroh S, Baxter C, Lessells R, Ogwell AE, Kebede Y, Tessema SK, de Oliveira T. 2022. Urgent need for a non-discriminatory and non-stigmatizing nomenclature for monkeypox virus. PLoS Biol 20:e3001769.

29. Vivancos R, Anderson C, Blomquist P, Balasegaram S, Bell A, Bishop L, Brown CS, Chow Y, Edeghere O, Florence I, Logan S, Manley P, Crowe W, McAuley A, Shankar AG, Mora-Peris B, Paranthaman K, Prochazka M, Ryan C, Simons D, Vipond R, Byers C, Watkins NA, team UMIM, Welfare W, Whittaker E, Dewsnap C, Wilson A, Young Y, Chand M, Riley S, Hopkins S, Monkeypox Incident Management T. 2022. Community transmission of monkeypox in the United Kingdom, April to May 2022. Euro Surveill 27.

30. Orviz E, Negredo A, Ayerdi O, Vazquez A, Munoz-Gomez A, Monzon S, Clavo P, Zaballos A, Vera M, Sanchez P, Cabello N, Jimenez P, Perez-Garcia JA, Varona S, Del Romero J, Cuesta I, Delgado-Iribarren A, Torres M, Sagastagoitia I, Palacios G, Estrada V, Sanchez-Seco MP, Grupo Viruela del Simio Madrid CNMIHS. 2022. Monkeypox outbreak in Madrid (Spain): Clinical and virological aspects. J Infect 85:412–417.

31. Yinka-Ogunleye A, Aruna O, Ogoina D, Aworabhi N, Eteng W, Badaru S, Mohammed A, Agenyi J, Etebu EN, Numbere TW, Ndoreraho A, Nkunzimana E, Disu Y, Dalhat M, Nguku P, Mohammed A, Saleh M, McCollum A, Wilkins K, Faye O, Sall A, Happi C, Mba N, Ojo O, Ihekweazu C. 2018. Reemergence of Human Monkeypox in Nigeria, 2017. Emerg Infect Dis 24:1149–1151.

32. Vaughan A, Aarons E, Astbury J, Balasegaram S, Beadsworth M, Beck CR, Chand M, O’Connor C, Dunning J, Ghebrehewet S, Harper N, Howlett-Shipley R, Ihekweazu C, Jacobs M, Kaindama L, Katwa P, Khoo S, Lamb L, Mawdsley S, Morgan D, Palmer R, Phin N, Russell K, Said B, Simpson A, Vivancos R, Wade M, Walsh A, Wilburn J. 2018. Two cases of monkeypox imported to the United Kingdom, September 2018. Euro Surveill 23.

33. Sadeuh-Mba SA, Yonga MG, Els M, Batejat C, Eyangoh S, Caro V, Etoundi A, Carniel E, Njouom R. 2019. Monkeypox virus phylogenetic similarities between a human case detected in Cameroon in 2018 and the 2017-2018 outbreak in Nigeria. Infect Genet Evol 69:8–11.

34. Erez N, Achdout H, Milrot E, Schwartz Y, Wiener-Well Y, Paran N, Politi B, Tamir H, Israely T, Weiss S, Beth-Din A, Shifman O, Israeli O, Yitzhaki S, Shapira SC, Melamed S, Schwartz E. 2019. Diagnosis of Imported Monkeypox, Israel, 2018. Emerg Infect Dis 25:980–983.

35. Yinka-Ogunleye A, Aruna O, Dalhat M, Ogoina D, McCollum A, Disu Y, Mamadu I, Akinpelu A, Ahmad A, Burga J, Ndoreraho A, Nkunzimana E, Manneh L, Mohammed A, Adeoye O, Tom-Aba D, Silenou B, Ipadeola O, Saleh M, Adeyemo A, Nwadiutor I, Aworabhi N, Uke P, John D, Wakama P, Reynolds M, Mauldin MR, Doty J, Wilkins K, Musa J, Khalakdina A, Adedeji A, Mba N, Ojo O, Krause G, Ihekweazu C, Team CDCMO. 2019. Outbreak of human monkeypox in Nigeria in 2017-18: a clinical and epidemiological report. Lancet Infect Dis 19:872–879.

36. Dumonteil E, Herrera C, Sabino-Santos G. 2023. Monkeypox Virus Evolution before 2022 Outbreak. Emerg Infect Dis 29:451–453.

37. Yong SEF, Ng OT, Ho ZJM, Mak TM, Marimuthu K, Vasoo S, Yeo TW, Ng YK, Cui L, Ferdous Z, Chia PY, Aw BJW, Manauis CM, Low CKK, Chan G, Peh X, Lim PL, Chow LPA, Chan M, Lee VJM, Lin RTP, Heng MKD, Leo YS. 2020. Imported Monkeypox, Singapore. Emerg Infect Dis 26:1826–1830.

38. CDC U. https://www.cdc.gov/poxvirus/mpox/response/2022/index.html. Accessed

39. WHO. https://worldhealthorg.shinyapps.io/mpx_global/. Accessed

40. Ruis C, Lusamaki E, O’Toole A, Otieno JR, Colquhoun R, Roemer C, Wawina-Bokalanga T, Tshiani-Mbaya O, Jansen D, Makangara JC, Agrawal A, Abu-Raddad LJ, Ahuka-Mundeke S, Ayouba A, Azhar EI, Caly L, Drosten C, Faria NR, Fowotade A, Hoxha A, Huang B, Korber B, le Polain de Waroux O, Lewis RF, Liesenborghs L, Melhem NM, Munster VJ, Ogoina D, Oude Munnink BB, Peiris JSM, Palacios G, Peeters M, Resende P, Rimoin AW, Saha S, Leo YS, Suzuki T, Vercauteren K, Yadav P, Van Kerkhove MD, Neher R, de Oliveira T, Ulaeto D, Mbala-Kingebeni P, Kindrachuk J, Koopmans MPG, Chand M, von Gottberg A, Subissi L, Rambaut A. 2025. A systematic nomenclature for mpox viruses causing outbreaks with sustained human-to-human transmission. Nat Med doi:10.1038/s41591-025-03820-6.

41. Masirika LM, Udahemuka JC, Schuele L, Nieuwenhuijse DF, Ndishimye P, Boter M, Mbiribindi JB, Kacita C, Lang T, Gortazar C, Musabyimana JP, Otani S, Aarestrup FM, Siangoli FB, Oude Munnink BB, Koopmans M. 2025. Epidemiological and genomic evolution of the ongoing outbreak of clade Ib mpox virus in the eastern Democratic Republic of the Congo. Nat Med 31:1459–1463.

42. Masirika LM, Udahemuka JC, Schuele L, Ndishimye P, Otani S, Mbiribindi JB, Marekani JM, Mambo LM, Bubala NM, Boter M, Nieuwenhuijse DF, Lang T, Kalalizi EB, Musabyimana JP, Aarestrup FM, Koopmans M, Oude Munnink BB, Siangoli FB. 2024. Ongoing mpox outbreak in Kamituga, South Kivu province, associated with monkeypox virus of a novel Clade I sub-lineage, Democratic Republic of the Congo, 2024. Euro Surveill 29.

43. Kinganda-Lusamaki E, Amuri-Aziza A, Fernandez-Nunez N, Makangara-Cigolo JC, Pratt C, Vakaniaki EH, Hoff NA, Luakanda-Ndelemo G, Akil-Bandali P, Nundu SS, Mulopo-Mukanya N, Ngimba M, Modadra-Madakpa B, Diavita R, Paku-Tshambu P, Pukuta-Simbu E, Merritt S, O’Toole A, Low N, Nkuba-Ndaye A, Kavunga-Membo H, Shongo Lushima R, Liesenborghs L, Wawina-Bokalanga T, Vercauteren K, Mukadi-Bamuleka D, Subissi L, Muyembe-Tamfum JJ, Kindrachuk J, Ayouba A, Rambaut A, Delaporte E, Tessema S, D’Ortenzio E, Rimoin AW, Hensley LE, Mbala-Kingebeni P, Peeters M, Ahuka-Mundeke S. 2025. Clade I mpox virus genomic diversity in the Democratic Republic of the Congo, 2018-2024: Predominance of zoonotic transmission. Cell 188:4–14 e6.

44. Vakaniaki EH, Kacita C, Kinganda-Lusamaki E, O’Toole A, Wawina-Bokalanga T, Mukadi-Bamuleka D, Amuri-Aziza A, Malyamungu-Bubala N, Mweshi-Kumbana F, Mutimbwa-Mambo L, Belesi-Siangoli F, Mujula Y, Parker E, Muswamba-Kayembe PC, Nundu SS, Lushima RS, Makangara-Cigolo JC, Mulopo-Mukanya N, Pukuta-Simbu E, Akil-Bandali P, Kavunga H, Abdramane O, Brosius I, Bangwen E, Vercauteren K, Sam-Agudu NA, Mills EJ, Tshiani-Mbaya O, Hoff NA, Rimoin AW, Hensley LE, Kindrachuk J, Baxter C, de Oliveira T, Ayouba A, Peeters M, Delaporte E, Ahuka-Mundeke S, Mohr EL, Sullivan NJ, Muyembe-Tamfum JJ, Nachega JB, Rambaut A, Liesenborghs L, Mbala-Kingebeni P. 2024. Sustained human outbreak of a new MPXV clade I lineage in eastern Democratic Republic of the Congo. Nat Med 30:2791–2795.

45. Americo JL, Earl PL, Moss B. 2023. Virulence differences of mpox (monkeypox) virus clades I, IIa, and IIb.1 in a small animal model. Proc Natl Acad Sci U S A 120:e2220415120.

46. Marennikova SS, Seluhina EM. 1976. Susceptibility of some rodent species to monkeypox virus, and course of the infection. Bull World Health Organ 53:13–20.

47. Osorio JE, Iams KP, Meteyer CU, Rocke TE. 2009. Comparison of monkeypox viruses pathogenesis in mice by in vivo imaging. PLoS One 4:e6592.

48. Stabenow J, Buller RM, Schriewer J, West C, Sagartz JE, Parker S. 2010. A mouse model of lethal infection for evaluating prophylactics and therapeutics against Monkeypox virus. J Virol 84:3909–20.

49. Earl PL, Americo JL, Moss B. 2012. Lethal monkeypox virus infection of CAST/EiJ mice is associated with a deficient gamma interferon response. J Virol 86:9105–12.

50. Earl PL, Americo JL, Moss B. 2017. Insufficient Innate Immunity Contributes to the Susceptibility of the Castaneous Mouse to Orthopoxvirus Infection. J Virol 91.

51. Earl PL, Americo JL, Cotter CA, Moss B. 2015. Comparative live bioluminescence imaging of monkeypox virus dissemination in a wild-derived inbred mouse (Mus musculus castaneus) and outbred African dormouse (Graphiurus kelleni). Virology 475:150–8.

52. Gigante CM, Korber B, Seabolt MH, Wilkins K, Davidson W, Rao AK, Zhao H, Smith TG, Hughes CM, Minhaj F, Waltenburg MA, Theiler J, Smole S, Gallagher GR, Blythe D, Myers R, Schulte J, Stringer J, Lee P, Mendoza RM, Griffin-Thomas LA, Crain J, Murray J, Atkinson A, Gonzalez AH, Nash J, Batra D, Damon I, McQuiston J, Hutson CL, McCollum AM, Li Y. 2022. Multiple lineages of monkeypox virus detected in the United States, 2021-2022. Science 378:560–565.

53. Hutson CL, Carroll DS, Gallardo-Romero N, Weiss S, Clemmons C, Hughes CM, Salzer JS, Olson VA, Abel J, Karem KL, Damon IK. 2011. Monkeypox disease transmission in an experimental setting: prairie dog animal model. PLoS One 6:e28295.

54. Hutson CL, Carroll DS, Self J, Weiss S, Hughes CM, Braden Z, Olson VA, Smith SK, Karem KL, Regnery RL, Damon IK. 2010. Dosage comparison of Congo Basin and West African strains of monkeypox virus using a prairie dog animal model of systemic orthopoxvirus disease. Virology 402:72–82.

55. Hutson CL, Gallardo-Romero N, Carroll DS, Clemmons C, Salzer JS, Nagy T, Hughes CM, Olson VA, Karem KL, Damon IK. 2013. Transmissibility of the monkeypox virus clades via respiratory transmission: investigation using the prairie dog-monkeypox virus challenge system. PLoS One 8:e55488.

56. Hutson CL, Olson VA, Carroll DS, Abel JA, Hughes CM, Braden ZH, Weiss S, Self J, Osorio JE, Hudson PN, Dillon M, Karem KL, Damon IK, Regnery RL. 2009. A prairie dog animal model of systemic orthopoxvirus disease using West African and Congo Basin strains of monkeypox virus. J Gen Virol 90:323–333.

57. Keckler MS, Carroll DS, Gallardo-Romero NF, Lash RR, Salzer JS, Weiss SL, Patel N, Clemmons CJ, Smith SK, Hutson CL, Karem KL, Damon IK. 2011. Establishment of the black-tailed prairie dog (Cynomys ludovicianus) as a novel animal model for comparing smallpox vaccines administered preexposure in both high- and low-dose monkeypox virus challenges. J Virol 85:7683–98.

58. Marennikova SS, Shelukhina EM, Zhukova OA. 1989. Experimental infection of squirrels Sciurus vulgaris by monkey pox virus. Acta Virol 33:399.

59. Saijo M, Ami Y, Suzaki Y, Nagata N, Iwata N, Hasegawa H, Iizuka I, Shiota T, Sakai K, Ogata M, Fukushi S, Mizutani T, Sata T, Kurata T, Kurane I, Morikawa S. 2009. Virulence and pathophysiology of the Congo Basin and West African strains of monkeypox virus in non-human primates. J Gen Virol 90:2266–71.

60. Xiao SY, Sbrana E, Watts DM, Siirin M, da Rosa AP, Tesh RB. 2005. Experimental infection of prairie dogs with monkeypox virus. Emerg Infect Dis 11:539–45.

61. Kilgore N, Nuzum EO. 2012. An interagency collaboration to facilitate development of filovirus medical countermeasures. Viruses 4:2312–6.

62. Seet BT, Johnston JB, Brunetti CR, Barrett JW, Everett H, Cameron C, Sypula J, Nazarian SH, Lucas A, McFadden G. 2003. Poxviruses and immune evasion. Annu Rev Immunol 21:377–423.

63. McInnes CJ, Damon IK, Smith GL, McFadden G, Isaacs SN, Roper RL, Evans DH, Damaso CR, Carulei O, Wise LM, Lefkowitz EJ. 2023. ICTV Virus Taxonomy Profile: Poxviridae 2023. J Gen Virol 104.

64. Liszewski MK, Leung MK, Hauhart R, Fang CJ, Bertram P, Atkinson JP. 2009. Smallpox inhibitor of complement enzymes (SPICE): dissecting functional sites and abrogating activity. J Immunol 183:3150–9.

65. Yadav VN, Pyaram K, Mullick J, Sahu A. 2008. Identification of hot spots in the variola virus complement inhibitor (SPICE) for human complement regulation. J Virol 82:3283–94.

66. Isaacs SN, Argyropoulos E, Sfyroera G, Mohammad S, Lambris JD. 2003. Restoration of complement-enhanced neutralization of vaccinia virus virions by novel monoclonal antibodies raised against the vaccinia virus complement control protein. J Virol 77:8256–62.

67. Rosengard AM, Liu Y, Nie Z, Jimenez R. 2002. Variola virus immune evasion design: expression of a highly efficient inhibitor of human complement. Proc Natl Acad Sci U S A 99:8808–13.

68. Spriggs MK, Hruby DE, Maliszewski CR, Pickup DJ, Sims JE, Buller RM, VanSlyke J. 1992. Vaccinia and cowpox viruses encode a novel secreted interleukin-1-binding protein. Cell 71:145–52.

69. Campbell JA, Trossman DS, Yokoyama WM, Carayannopoulos LN. 2007. Zoonotic orthopoxviruses encode a high-affinity antagonist of NKG2D. J Exp Med 204:1311–7.

70. Maluquer de Motes C, Cooray S, Ren H, Almeida GM, McGourty K, Bahar MW, Stuart DI, Grimes JM, Graham SC, Smith GL. 2011. Inhibition of apoptosis and NF-kappaB activation by vaccinia protein N1 occur via distinct binding surfaces and make different contributions to virulence. PLoS Pathog 7:e1002430.

71. Cooray S, Bahar MW, Abrescia NGA, McVey CE, Bartlett NW, Chen RA, Stuart DI, Grimes JM, Smith GL. 2007. Functional and structural studies of the vaccinia virus virulence factor N1 reveal a Bcl-2-like anti-apoptotic protein. J Gen Virol 88:1656–1666.

72. Katoh K, Standley DM. 2013. MAFFT multiple sequence alignment software version 7: improvements in performance and usability. Mol Biol Evol 30:772–80.

73. Guindon S, Gascuel O. 2003. A simple, fast, and accurate algorithm to estimate large phylogenies by maximum likelihood. Syst Biol 52:696–704.

